# Abbapolin inhibitors of the PLK1 PBD as Prostate Cancer Therapeutics, in vivo activity and synergy with androgen therapy

**DOI:** 10.64898/2026.07.02.736204

**Authors:** George Merhej, Gurusankar Ramamoorthy, Danda Chapagai, Mohammad Esfini Farahani, Yifan Kong, Chintada Nageswara Rao, Jessy Stafford, Zachary T. Mack, Cassidy Socia, Shikha Kumari, Kristen Hogan, Niti Jani, Maria Marjorette Pena, Elmar Nurmemmedov, Ivan Babic, Mengqian Chen, Xiaoqi Liu, Michael D. Wyatt, Campbell McInnes

## Abstract

Polo-like kinase 1 (PLK1) is an established therapeutic target in cancer; however, ATP-competitive kinase inhibitors have shown limited clinical success because of toxicity, acquired resistance, and incomplete inhibition of non-catalytic PLK1 functions. Targeting the Polo-box domain (PBD), which regulates PLK1 localization and substrate recognition, represents an alternative therapeutic strategy but has been hindered by the lack of selective, cell-active small molecules. Here, the optimization and biological characterization of abbapolins, a series of non-peptidic PLK1 PBD inhibitors developed using the REPLACE strategy are described. Structure-guided optimization and screening across the NCI-60 cancer cell panel identified compounds with preferential activity against prostate cancer cells. Proteomic analyses demonstrated that cellular sensitivity correlated with PLK1 protein abundance, supporting an on-target mechanism of action. Abbapolins directly engaged PLK1 in cells, induced selective degradation of endogenous PLK1, and suppressed long-term clonogenic growth. Lead compounds demonstrated favorable pharmacokinetic properties and significantly inhibited prostate tumor growth in xenograft models without detectable systemic toxicity. PLK1 abundance was significantly reduced in treated tumors and correlated with tumor response, identifying PLK1 degradation as a potential pharmacodynamic biomarker. Abbapolins also synergized with enzalutamide in castration-resistant prostate cancer cells, supporting their potential as combination therapies for advanced disease. Collectively, these studies establish selective inhibition of the PLK1 Polo-box domain as a viable therapeutic strategy, provide in vivo proof-of-concept for the REPLACE approach, and identify abbapolins as promising leads for advanced prostate cancer.

## INTRODUCTION

Polo Like Kinases (PLKs) are Serine/Threonine kinases that regulate cell cycle progression[1, 2] with the individual enzymes (PLK1-5) having unique expression, roles, and functions, substrates, and sub-cellular location[3–5]. PLK1-3 have a conserved N-terminal kinase domain (KD) and a C-terminal polo box domain (PBD) but diverge significantly in the interdomain connecting loop (IDL), which varies in length and composition[3]. As the IDL is known to affect the activity of PLK1, it likely plays a role in the differential regulation of other PLK members. Furthermore, the residues stabilizing the closed confirmation in PLK1 diverge in the other PLKs, leading to varying conformation regulation[3]. PLK1 is the most extensively characterized, leading to identification of its critical roles regulating entry into and progression through mitosis (centrosome maturation, bipolar spindle formation, sister chromatin separation, and cytokinesis)[6, 7] and is known to localize at key mitotic structures including centrosomes and kinetochores among others[8]. The PBD, which binds a phosphoserine/phosphothreonine (STP) motif on target proteins, is critical for its sub-cellular localization and for substrate recognition prior to phosphorylating target substrates[9–12].

PLK1 is highly expressed in many tumor types and levels are prognostic of unfavorable outcomes[13–15]. PLK1 has been extensively investigated as a drug target both genetically via specific mutations/deletions of tumor-driving genes and through development of ATP-competitive compounds that sensitize tumor cells dependent on PLK1[16–18]. The latter approach has resulted in several compounds advancing to clinical trials, including volasertib and onvansertib, although clinical progress has yet to achieve US FDA approval. Specificity for PLK1 versus PLK2 and PLK3 is important as these family members have opposing functions and thus act as tumor suppressors[19]. A single point mutation (C67V) in the KD confers dramatic cellular resistance *in vitro* to several KD inhibitors [20]. While PLK1 mutations conferring resistance to KD inhibitors are not yet clinically observed, point mutations lead to significant resistance issues in other kinases targeted in oncology. Another potentially deleterious, unintended consequence identified in recent studies is that KD inhibitors induce an open conformation of PLK1[4, 21]. This conformation, while catalytically inhibited, has increased affinity for peptides from key substrates [21, 22]. In other words, KD inhibitors may not block non-catalytic roles of PLK1 and in fact may promote non-catalytic PBD functions of PLK1.

The potential drawbacks of KD inhibitors justify the development of PLK1 inhibitors through alternate approaches[23, 24]. The phospho-recognition motifs in the PBD are unique to the PLK family, and peptides and small molecules have been demonstrated to bind selectively to the PLK1 PBD[24]. HeLa cells overexpressing the PBD of PLK1 arrested in mitosis and possessed chromosome congression defects[8], suggesting that stoichiometry and regulation of the PBD itself are important. Even though the PBD phosphorecognition site is conserved among PLKs, phosphopeptides bind preferentially to the PLK1 PBD and therefore its targeting provides an opportunity for blocking PLK1 PBD functions selectively. Several of the small molecule PBD inhibitors [25–28] identified thus far turned out to be non-specific protein alkylators that have little or no potential to be PLK1 PBD-specific inhibitors[28]. While other PBD inhibitors have been reported, no proof of direct engagement of PLK1 in cells exists [25, 26, 29].

REPLACE (Replacement with Partial Ligand Alternatives through Computational Enrichment) is a validated strategy for the conversion of peptide inhibitors to generate more drug-like leads and has been used to discover novel PBD inhibitors in an iterative fashion. The compounds, named abbapolins, exhibit good affinity for the PLK1 PBD, engage and lead to degradation of endogenous PLK1, and show potent anti-proliferative activity against tumor cells, as well activity in cells expressing C67V PLK1 that are dramatically resistant to KD-targeted compounds [30, 31]. Furthermore, abbapolins have been shown to block dimerization of PLK1, a key regulatory step in G2 and mitosis[21]. Here, further development of the abbapolin series has been undertaken through biochemical, cellular engagement, and antiproliferative assays. Evaluation of analogs in the NCI-60 cell line panel allowed their cellular structure-activity relationship (SAR) to be established. Following up from the panel results with detailed studies in prostate cancer cell-lines including clonogenic assays revealed highly potent antiproliferative activity. Experiments with androgen-resistant prostate cancers demonstrated a synergistic effect and sensitization to enzalutamide treatment. This cellular efficacy then triggered evaluation in *in vivo* models to determine pharmacokinetic (PK) and pharmacodynamic (PD) activity. The results demonstrated blood levels approaching the cellular IC_50_ of the compounds and a statistically significant tumor growth inhibition in a xenograft model without observable gross toxicity, which confirms the potential of abbapolins as potential therapeutics for hormone-refractory prostate cancer. Overall, results obtained provide *in vivo* proof-of-concept for the REPLACE strategy and for targeting the PBD as an alternative strategy to inhibiting the KD of PLK1.

## METHODS

### Chemical Synthesis General Methods

Unless stated otherwise, all reactions were carried out in oven or flame-dried glassware under an atmosphere of dry nitrogen. All reaction mixtures were stirred magnetically. A silicon oil bath was used as the heat source for reactions performed above room temperature. Air and moisture-sensitive liquids were transferred via syringe using a standard technique. ^1^H and ^13^C nuclear magnetic resonance (NMR) spectra were recorded in CDCl_3_ unless otherwise mentioned in MeOH-d_4_ or DMSO-d_6_. NMR spectra were recorded using a Bruker 300 or 400 MHz. Chemical shifts (*δ* H) are reported in parts per million (ppm) relative to the residual solvent signal of CDCl_3_ (0.00 ppm; TMS signal), CDCl_3_ (7.16 ppm), or DMSO-*d_6_* (2.50 ppm). ^1^H-NMR coupling constants (J) are reported in Hertz (Hz) and refer to apparent multiplicities. Data are reported as follows: chemical shift, multiplicity (s = singlet, br s = broad singlet, d = doublet, t = triplet, q = quartet, quin = quintet, sext = sextet, sept = septet, m = multiplet, dd = doublet of doublets, etc.), coupling constant, and integration. ^13^C-NMR spectra were recorded at either 75 or 101 MHz. Chemical shifts (*δ* C) are reported in ppm relative to CDCl_3_ (77.16 ppm), or DMSO-*d_6_* (39.57 ppm). Analytical thin-layer chromatography (TLC) was performed on Silica Gel 60 F254 Coated Aluminum-Backed TLC plates (60 Å porosity, F-254 indicator). Unless stated otherwise, compounds were visualized by UV irradiation. All commercial solvents were purchased from Fisher or VWR unless otherwise mentioned and used without further distillation. Deionized water was used for aqueous reactions. All other reagents were purchased from various commercial sources and used as received without further purification. For detailed synthetic methods and characterization please see the supplementary information.

### Cell Culture, Colony Formation, and Immunoblotting

PC3 prostate cancer cells (ATCC, Manassas, VA, USA) were cultured in Ham’s F-12K medium (Cellgro, Manassas, VA, USA) supplemented with 10% Nu-serum and 1% penicillin-streptomycin. Two thousand PC3 cells were seeded into 6-well plates for 24 hours and treated with drug concentrations described in the figure legends. After 10 days, the colonies were fixed with methanol and stained with Giemsa (10%). Image J was used to count colonies. Means and standard deviations were calculated from three independent experiments. Protein expression analysis was performed via immunoblotting, as previously described[21, 31]. For protein analysis, in brief, protein concentrations were determined using the BCA protein assay kit (#23225, Pierce, Rockford, IL, USA). Equal amounts of protein (25 µg) were resolved on 4–15% SDS-PAGE gels. The following primary antibodies were used: anti-PLK1(#05-844, Millipore Sigma, Temecula, CA, USA), anti-phospho-TCTP (Ser 46) (#5251, Cell Signaling Technology, Danvers, MA, USA), and anti-GAPDH (#5174, Cell Signaling Technology). Detection was carried out using HRP- conjugated secondary antibodies (#NA931 and #NA934, GE Healthcare, Little Chalfont, Buckinghamshire, UK) and a chemiluminescence detection kit (#32106, ThermoFisher, Waltham, MA, USA), following the manufacturer’s protocol. For tumor protein analysis, tumors were dissected on ice and 5 mg of each tumor tissue was used for protein extraction. The tumor tissue was snap-frozen by immersion in liquid nitrogen and 500 µL RIPA lysis buffer (1% protease-phosphatase inhibitor cocktail) was then added. The tumor tissue was then homogenized on ice using an electric homogenizer, then samples were agitated in lysis buffer for 2 hours at 4° C. The samples were centrifuged at 16,000 x g for 20 minutes at 4° C. The supernatants were collected and stored at -80° C. Quantification of protein concentration and immunoblotting were performed in a similar manner to cellular protein extracts.

### Androgen Resistance and Combination Assays

C4-2R castration-resistant prostate cancer cells were kindly provided by Dr. Allen Gao (University of California, Davis, CA, USA) and maintained in RPMI-1640 medium supplemented with 10% fetal bovine serum (FBS) and 1% penicillin-streptomycin at 37°C in a humidified atmosphere of 5% CO₂. For colony formation assays, two thousand C4-2R cells were seeded in 6-well plates and allowed to adhere overnight. Cells were then treated with enzalutamide alone (0, 10, 20, or 40 µM), abbapolin **6** alone (0, 10, 20, or 40 µM), or with both agents in combination across the full 4×4 dose matrix. The medium containing fresh drug was replaced every three days. After 10 days, colonies were fixed with 4% formaldehyde, stained with crystal violet (0.5% w/v), and counted. Colony counts were normalized to the untreated control to calculate percent survival.

For synergy analysis, C4-2R cells were seeded in 96-well plates and treated with enzalutamide (0, 10, 20, or 40 µM) and abbapolin **6** (0, 10, 20, or 40 µM) as single agents and in all pairwise combinations across a 4×4 dose matrix. Cell viability was assessed using the MTS assay (Cell Titer 96® Aqueous One Solution, Promega) according to the manufacturer’s instructions. Cell viability values, normalized to untreated control wells, were uploaded to Synergy Finder (version 3.0; https://synergyfinder.fimm.fi)[32], which calculated synergy scores across the dose matrix. Drug interaction was scored using the Loewe model. Synergy scores, heatmaps, and 3D landscape plots were generated within Synergy Finder to visualize the interaction across the tested concentration range.

### Tumor Xenograft Experiments

PC3 and 22RV1 human prostate carcinoma cells (CRL-2505, ATCC, Manassas VA USA) cells were grown as described above to ∼75% confluence, collected on the day of injection and resuspended in 50% matrigel. One million cells in 75 microliters were injected subcutaneously (s.c.) into the flank of 8- week old athymic nude mice (JAX 002019, Jackson Laboratory, Bar Harbor Maine, USA). Tumors were allowed to grow to 200 mm^3^, Mice were then randomly assigned to control (vehicle) or treatment groups (inhibitors). Vehicle or inhibitors were administered by intraperitoneal injection (i.p.) or by oral gavage (o.g.) five days a week for three weeks. Tumor volumes were measured daily using calipers, and volume was calculated as ((width)^2^ x length)/2). Tumors and organs were harvested at sacrifice and weights were recorded. For the initial 22Rv1 and PC3 experiments (supplementary figures 1 and 2), abbapolin **6** was administered at 35 mg/kg i.p. (dosing vehicle PBS) and **7** at 100 mg/kg o.g.;(dosing vehicle 30% propylene glycol and 70% PEG400). B16727 was administered o.g at 50 mg/kg two days per week. Control mice were given phosphate buffered saline (PBS). For the repeat PC3 experiment (Figure 5), abbapolin **6** was administered at 55 mg/kg i.p. (100 μL, dosing vehicle PBS) and **8** at 150 mg/kg o.g. (dosing vehicle 30% propylene glycol and 70% PEG400).

## RESULTS

### Structure-activity relationship and lead optimization of the abbapolins

The development of non-ATP competitive PLK1 inhibitors targeting the Polo-Box domain through the REPLACE strategy resulted in the identification of abbapolins. These non-peptidic molecules have confirmed cellular engagement leading to the degradation of endogenous PLK1 and show potent anti-proliferative activity against tumor cells[30, 31]. Here, further investigation of the abbapolin series is described through the development of novel structural analogs designed to better engage both the phospho-binding motif of the PBD as well as the hydrophobic cryptic pocket of PLK1. Previously described abbapolins were based solely upon mimicking the interactions of the substrate phosphothreonine with a carboxylate isostere. Substrates with priming phosphorylation on the PBD- engaging motif interact through ion pairing with H538 and K540 of the PBD. To further explore the structure-activity of this series, a library of compounds incorporating phosphate isosteres (**Table 1**) was generated, and these compounds were tested against PC3 prostate cancer cells in a 72-hour MTT assay. The obtained results were also compared to the carboxylate isosteres[31]. In the first instance, compound **1** the phosphate isostere of the carboxylate in the ortho position (R2, Table 1) was generated. Compounds **2** and **3** were found to be 2 to 3-fold more potent in the MTT assay than compounds with the same 8-atom alkyl tail [31]. Replacement of the R2 carboxylate with a phosphonate group instead of the phosphate led to an even greater cellular potency enhancement, with an IC_50_ of 9.2 μM (**3**). The phosphonate isostere directly connected to the aromatic ring was, however, significantly less potent (**4**) than both of these, but more so than the parent carboxylate. The analog containing a phosphonate at the meta-position (R3, Table 1) showed minimal activity, suggesting that the additional oxygen acts as a spacer to appropriately position the dianion for interaction with H538 and K540 of the PBD (**5**). Further SAR of the phosphorous isosteres showed that moving the phosphate to the meta position (R3) led to the most potent of this series with an IC_50_ of 2.3 μM (**6**). In addition to phosphate isosteres, an analog, (**7**) replacing the amide bond linkage with a sulfonamide was generated. This compound, which also contains a fluorine atom at R2, was significantly more potent with a 3-fold greater IC_50_ in PC3 cells compared to the parent amide derivative[31].

**Table 1.**
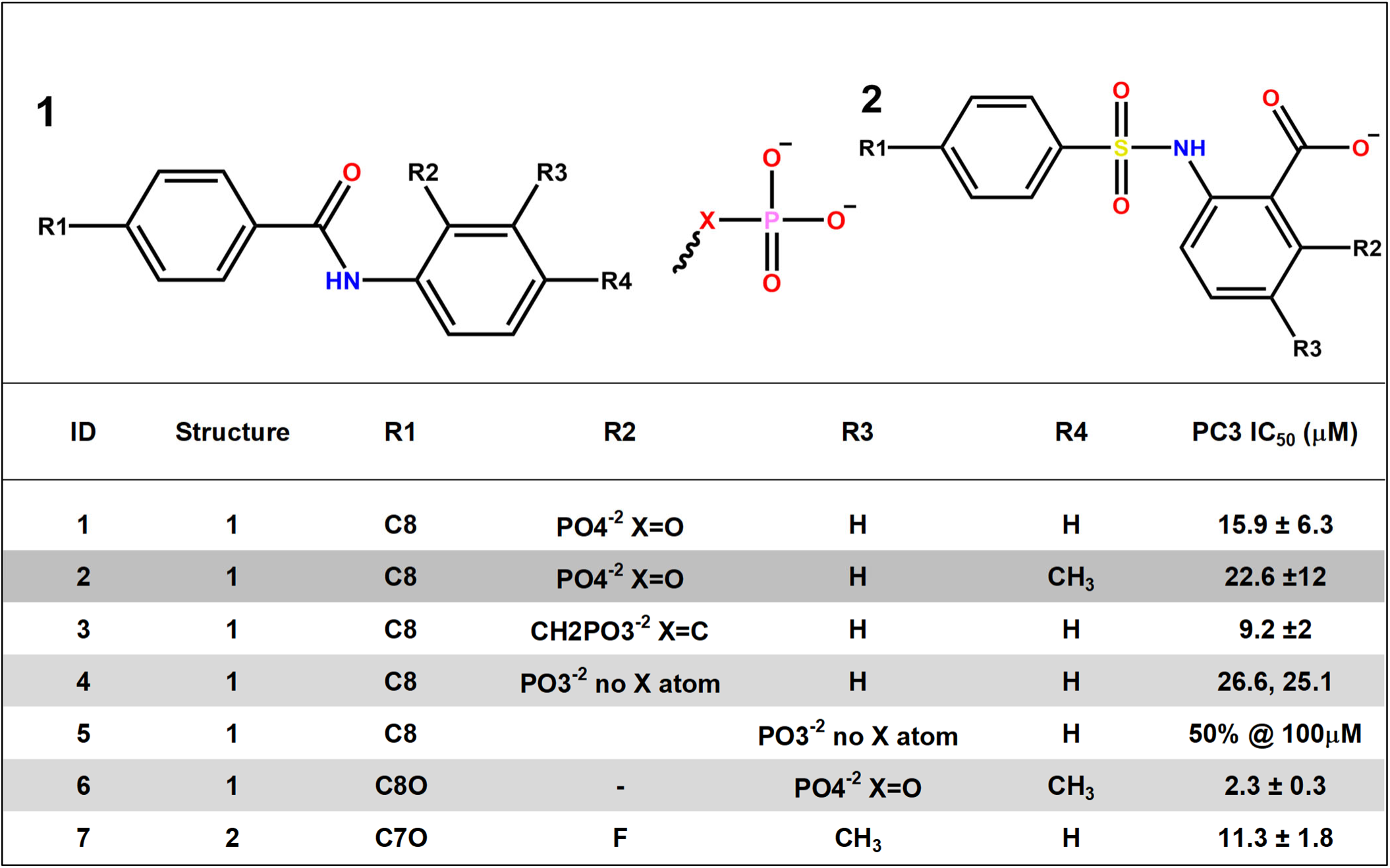
Structure-activity relationship of abbapolin phosphate isosteres in PC3 cells.

### NCI-60 Screening of abbapolins and further evaluation in colony-forming assays

To further explore the development potential of the abbapolins, compounds previously described as well as newly optimized analogs were evaluated in the NCI-60 panel, part of the National Cancer Institute (NCI) Developmental Therapeutics Program. In general, compounds are initially tested at a single dose in 60 cell lines derived from 9 different tissue types, and those that show reasonable activity in the majority of tumors are then put forward for dose-response testing in the panel. Analysis of results for abbapolins from the single dose and dose-response testing revealed some selective vulnerabilities, with prostate cancer (PC3 and DU145), leukemias, CNS tumors, and colorectal tumors responding the best to abbapolin treatment. The sensitive cell lines contrast with, for example, breast cancer cells for which abbapolins are 2-8 fold less active. Using the NCI-60 results for the best responding abbapolins after dose-response testing, a COMPARE analysis was carried out to explore patterns of activity in relation to publicly available data for molecules that have been screened in the panel. The COMPARE algorithm is a powerful tool developed by NCI to explore and interpret patterns of activity by comparing a compound’s antiproliferative fingerprint across the 60 cell lines to historical and contemporary data. COMPARE can therefore provide suggestions of potential common mechanisms of action, tumor selectivity, and identify phenotypic analogs. Initially, the APPROVEDDRUGS dataset was compared with the clinically evaluated PLK1 KD inhibitor in the database, volasertib, which is well known to induce mitotic arrest. Not surprisingly, the top comparator to volasertib was the antimitotic paclitaxel (taxol), with a Pearson correlation coefficient of 0.78. Note that Pearson values above 0.8 are considered strongly suggestive, above 0.7 are considered suggestive, and a value of 0.6 is considered the lowest value inferring any relatedness of the mechanism of action for two compounds. Four abbapolins for which dose response testing occurred were compared against the APPROVEDDRUGS set, **8, 9, 11**, and **14**. Because these compounds were tested by the DTP program prior to the conversion to the 384-well format, our compounds were analyzed against the “Classic Screen.” COMPARE revealed that there was no drug in the dataset with a Pearson correlation coefficient above 0.6 for any of the abbapolins, and only one single compound for one abbapolin above 0.5. These analyses suggest that the PBD-targeted abbapolins have a unique mechanism of action in relation to all of the FDA-approved drugs in COMPARE at the time the comparisons were run.

The abbapolins were also analyzed by COMPARE against the ALLNSC data set to compare their activities against all compounds tested in the NCI-60 screen to investigate potential commonalities in mechanism of action with all molecules available in the dataset. COMPARE of compounds **8, 9,** and **11** all revealed Pearson values of 0.85-0.91 for each other, strongly suggesting that they each possessed similar mechanisms of action. For example, the query with **8** revealed a Pearson value of 0.88 with **11** and 0.91 with **9**. A query with **9** revealed a Pearson value of 0.91 for compound **11**. Aside from each other, COMPARE revealed no other compounds with a Pearson value above 0.7, and only 2-4 compounds above 0.6, again suggesting a unique mechanism of action for these abbapolins with the aminoalkyl tails. None of these weakly correlating compounds were structurally similar to the abbapolins and are unlikely to be KD or PBD PLK1 inhibitors or provide useful mechanistic insights. When compound **14** with the sulfur heteroatom connecting the benzamide to the alkyl tail was queried, it was intriguing to find that the Pearson values for **8, 9,** and **11**, each showed values below 0.6 in relation to **14**. Indeed there were no compounds across the NSC dataset with Pearson values above 0.65 for **14** and only five compounds with values above 0.6, suggesting this analog possess a distinctive activity profile that remains uninvestigated at this time.

Through further analysis of the NCI-60 data set obtained, the structure-activity relationship for the features of the abbapolins that led to the best anti-proliferative activity was established and the results are summarized in **Table 2**. The most active compounds were those containing a thioether or amine group connecting the alkyl tail with the benzamido group and which engages the cryptic pocket in the PBD substrate recognition site. Firstly, the amino alkyl series consisting of compounds **8-11** demonstrated the broadest activity against the NCI-60 panel when tested at 10 μM. Of these, **8,** possessing a C7NH tail and methyl at R3, exhibited growth inhibition of 50% or more for 39 of the 60 cell lines in the panel, while **9** with a C8NH alkyl tail and methyl at R3 inhibited 37 of 60 cell lines at this dose. Compound **10**, the C8NH version of **8** surprisingly only inhibited 9 cell lines, while compound **11**, the N-methylated version of **9,** inhibited 25 lines. Of all compounds tested in the NCI-60 panel, the alkyamine derivatives **8**, **9** , and **11** exhibited the most potent activity against prostate tumors with low micromolar IC_50_ values being observed after dose response testing. For the thioether series, the most broadly potent compound was **14** with a C8S tail and an R2 methyl, inhibiting 42 or the 60 cell lines of 50% or greater.

**Table 2.**
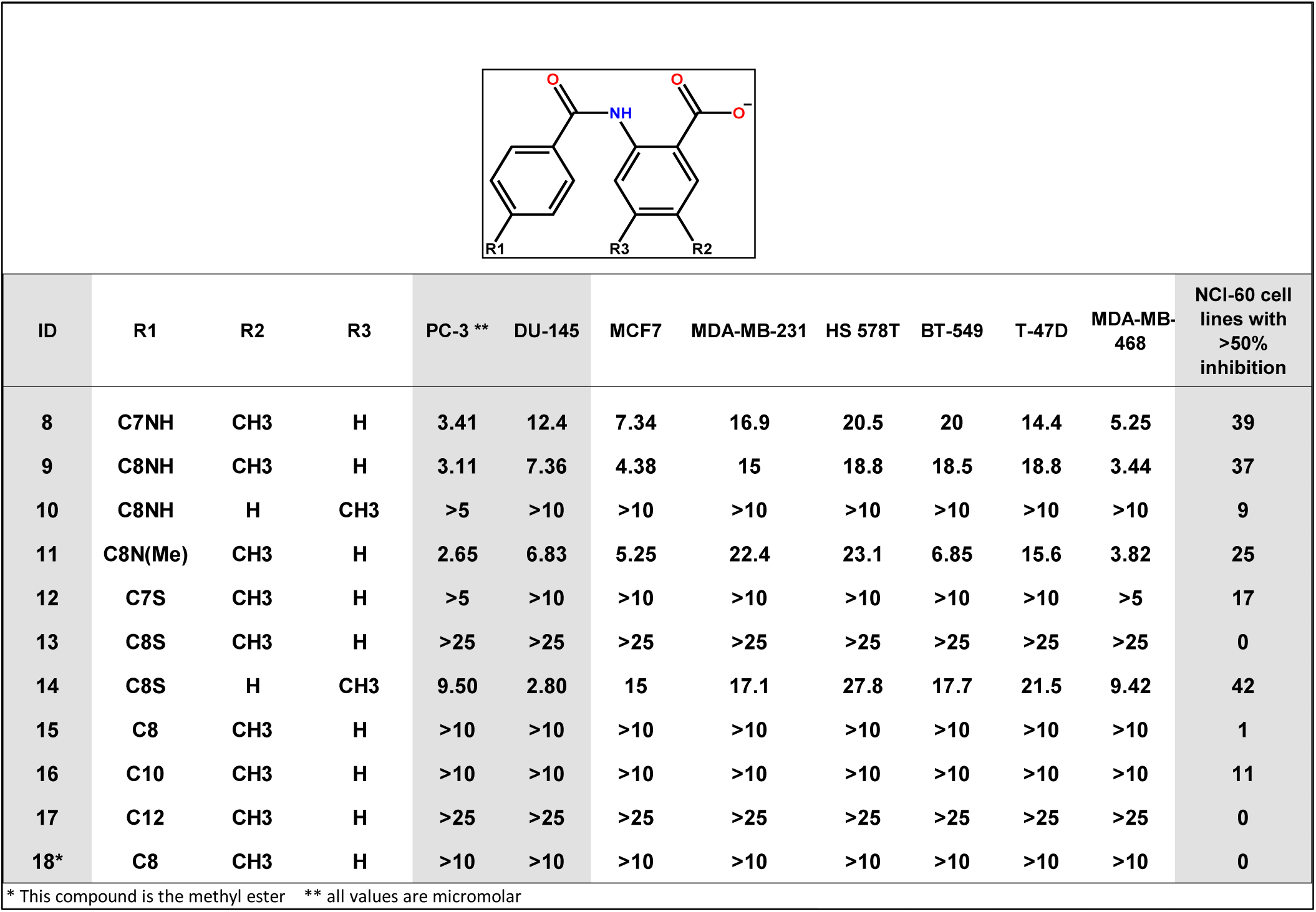
Structure-activity relationship on Prostate and Breast cancer cell lines in the NCI-60 panel.

| ID | R1 | R2 | R3 | PC-3 ** | DU-145 | MCF7 | MDA-MB-231 | HS 578T | BT-549 | T-47D | MDA-MB-468 | NCI-60 cell lines with >50% inhibition |
| --- | --- | --- | --- | --- | --- | --- | --- | --- | --- | --- | --- | --- |
| 8 | C7NH | CH3 | H | 3.41 | 12.4 | 7.34 | 16.9 | 20.5 | 20 | 14.4 | 5.25 | 39 |
| 9 | C8NH | CH3 | H | 3.11 | 7.36 | 4.38 | 15 | 18.8 | 18.5 | 18.8 | 3.44 | 37 |
| 10 | C8NH | H | CH3 | >5 | >10 | >10 | >10 | >10 | >10 | >10 | >10 | 9 |
| 11 | C8N(Me) | CH3 | H | 2.65 | 6.83 | 5.25 | 22.4 | 23.1 | 6.85 | 15.6 | 3.82 | 25 |
| 12 | C7S | CH3 | H | >5 | >10 | >10 | >10 | >10 | >10 | >10 | >5 | 17 |
| 13 | C8S | CH3 | H | >25 | >25 | >25 | >25 | >25 | >25 | >25 | >25 | 0 |
| 14 | C8S | H | CH3 | 9.50 | 2.80 | 15 | 17.1 | 27.8 | 17.7 | 21.5 | 9.42 | 42 |
| 15 | C8 | CH3 | H | >10 | >10 | >10 | >10 | >10 | >10 | >10 | >10 | 1 |
| 16 | C10 | CH3 | H | >10 | >10 | >10 | >10 | >10 | >10 | >10 | >10 | 11 |
| 17 | C12 | CH3 | H | >25 | >25 | >25 | >25 | >25 | >25 | >25 | >25 | 0 |
| 18* | C8 | CH3 | H | >10 | >10 | >10 | >10 | >10 | >10 | >10 | >10 | 0 |

\* This compound is the methyl ester \*\* all values are micromolar

Compounds **12** and **13** were less potent and **17** showed no activity in any cancer cell line tested. For the compounds without a heteroatom and those possessing an alkyl chain, lower activities, including C8 (**15**), C10 (**16**) and C12 (**17**) were observed, with only compound **16** showing a significant number of cell lines inhibited.

Although the data from the NCI-60 screening was encouraging and in broad concurrence with published work [30, 31] from this laboratory, it was noted that several of the compounds had lower than expected activity in the NCI-60 panel based on prior analyses. This included all of the phosphate isosteres which, despite having significant activity in PC3 cells in the 72-hour MTT assay, showed little activity in the NCI-60 panel. In particular, **6**, which had low μM IC_50_ values in PC3 cells in-house, showed little inhibition in the PC3 and DU145 lines in the NCI-60 panel. Compounds **6** and **8** were therefore further investigated using colony forming assays to assess longer-term exposure in contrast to the MTT assay (72 h) and NCI-60 screen (48 h) that utilize shorter drug exposure times. The results of colony-forming assays showed that **6** inhibited colony formation of PC3 cells with an IC_50_ of 0.337 ± 0.045 μM, and **8** inhibited growth with an IC_50_ of 6.28 ± 1.05 μM, suggesting that for the phosphate analogs the shorter-term assays were missing a feature of the onset of action.

### Proteomic analysis of abbapolins and correlation with PLK1 and substrate levels

To determine whether total PLK1 protein correlates with the phosphoforms pT210 and pT214, two critical phospho-sites for PLK1 function[33], correlation relation of mass spectrometry-based proteomics was performed utilizing ATLANTiC as reported[34]. Total PLK1 protein was significantly correlated with pT214 PLK1 levels (Pearson’s r = 0.57, P = 0.0002, n = 38 cell lines) (**Figure 3A**). However, no significant correlation was observed between total PLK1 and pT210 PLK1, which might be due to an inability to detect this form which is specific for the G2/M boundary and thus undetectable in an asynchronous cell population. Whether the antiproliferative activity of compound **8** is associated with PLK1 expression and its phosphorylated states was then investigated. Correlation analyses were performed between the IC_50_values of compound **8** and the levels of total and phosphorylated PLK1 across the NCI-60 panel. The IC_50_ values of compound **8** showed a significant inverse correlation with total PLK1 protein (Pearson’s r = −0.32, P = 0.01, n = 60 cell lines) (**Figure 3B**), suggesting that cells expressing higher levels of PLK1 are more sensitive to compound **8.** These findings further support an on-target mechanism of action for the compound. However, no significant correlation was observed between the IC_50_ values of compound **8** and the phosphorylated forms of PLK1 at T210 or T214. This may be explained by the fact that phosphorylation at T210 and T214 is tightly regulated for entry into mitosis, whereas the activity of compound **8** acts outside of the cell cycle phase in which these phosphoforms are most abundant. It was previously observed that in contrast to KD inhibitors, abbapolins do not induce a mitotic arrest, suggesting they interfere with PLK1 prior to mitosis[31].

**Figure 1.**
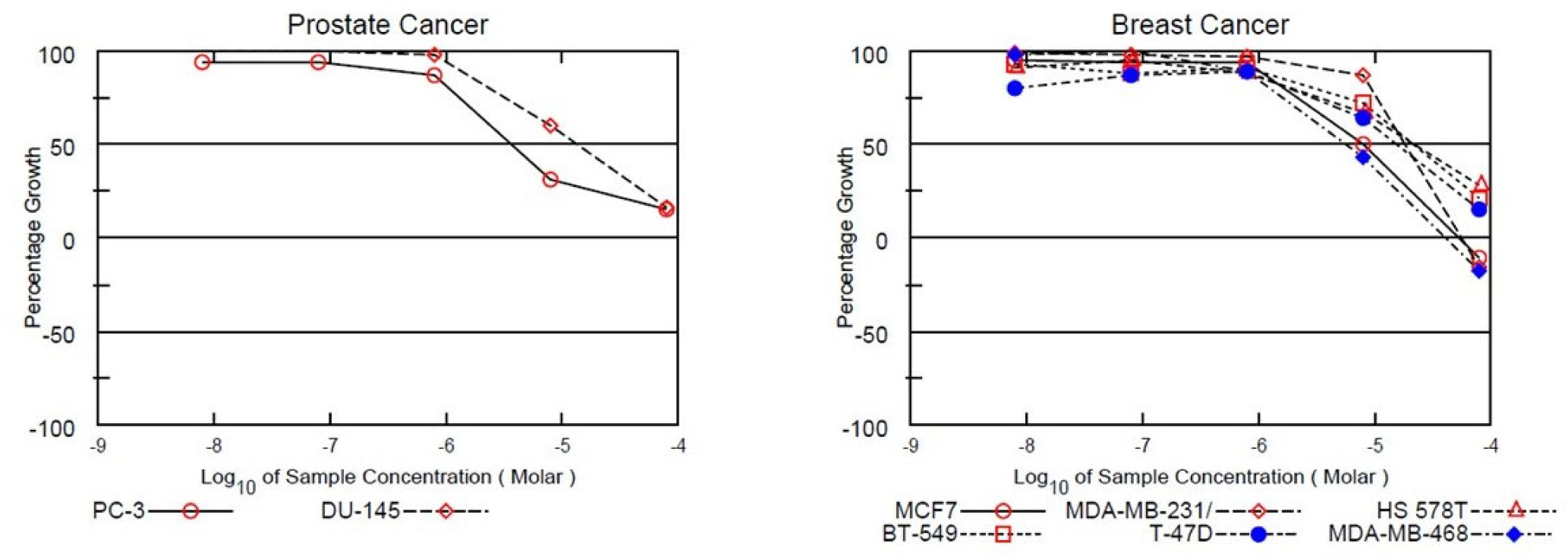
Subset of results from the NCI-60 tumor cell line panel. Compound **8** (Table 2) showed broad activity in initial single dose screening and was then evaluated in a dose response testing. Prostate cancers were among the most sensitive with breast tumors being less so.

**Figure 2.**
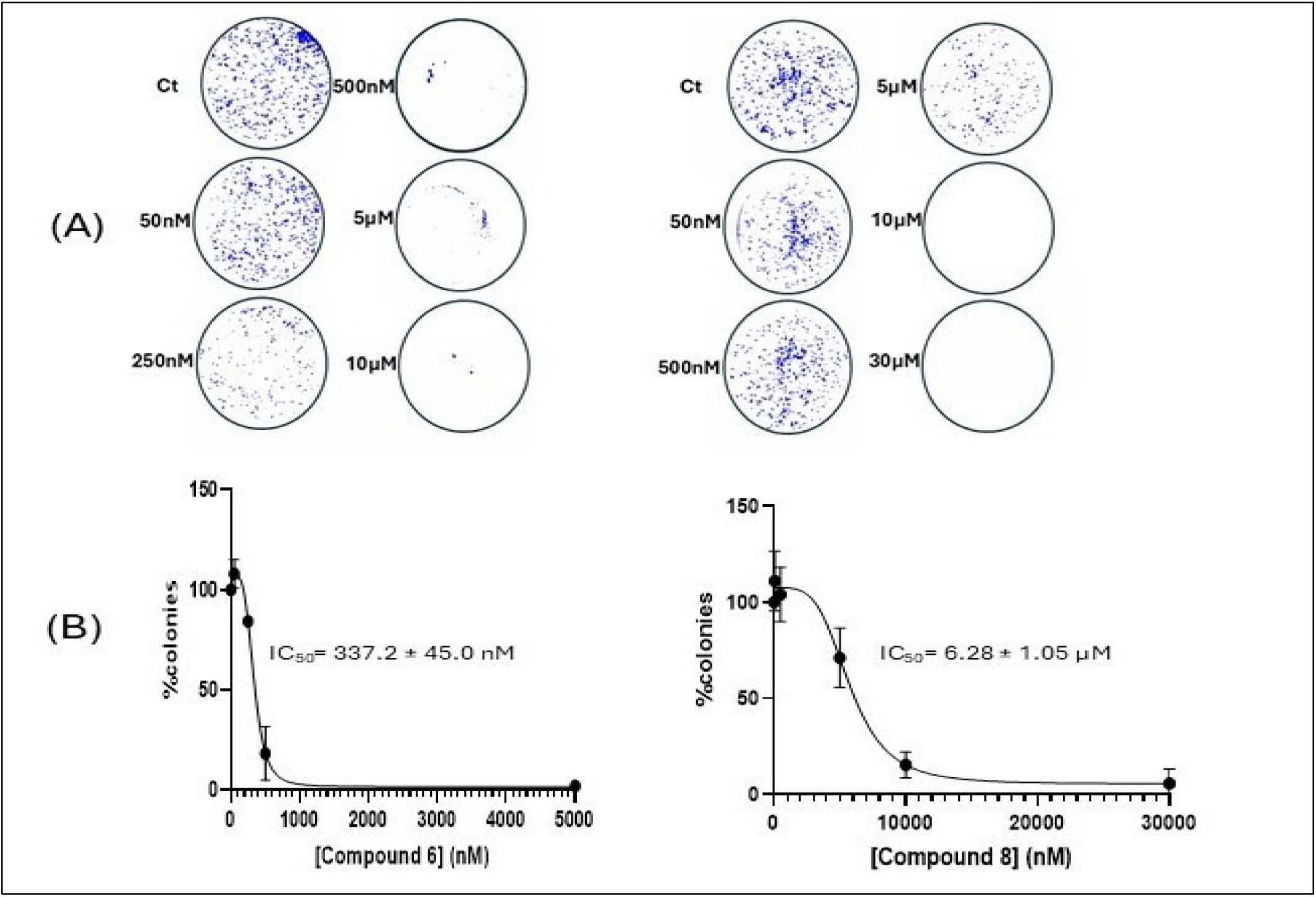
Abbapolin activity in Colony Forming Assays. PC3 cells were treated with compounds **6** (A and B left) and **8** (A and B right) for 10 days, and colony formation was assessed. The data are the mean ± SD (n=3 independent experiments).

**Figure 3.**
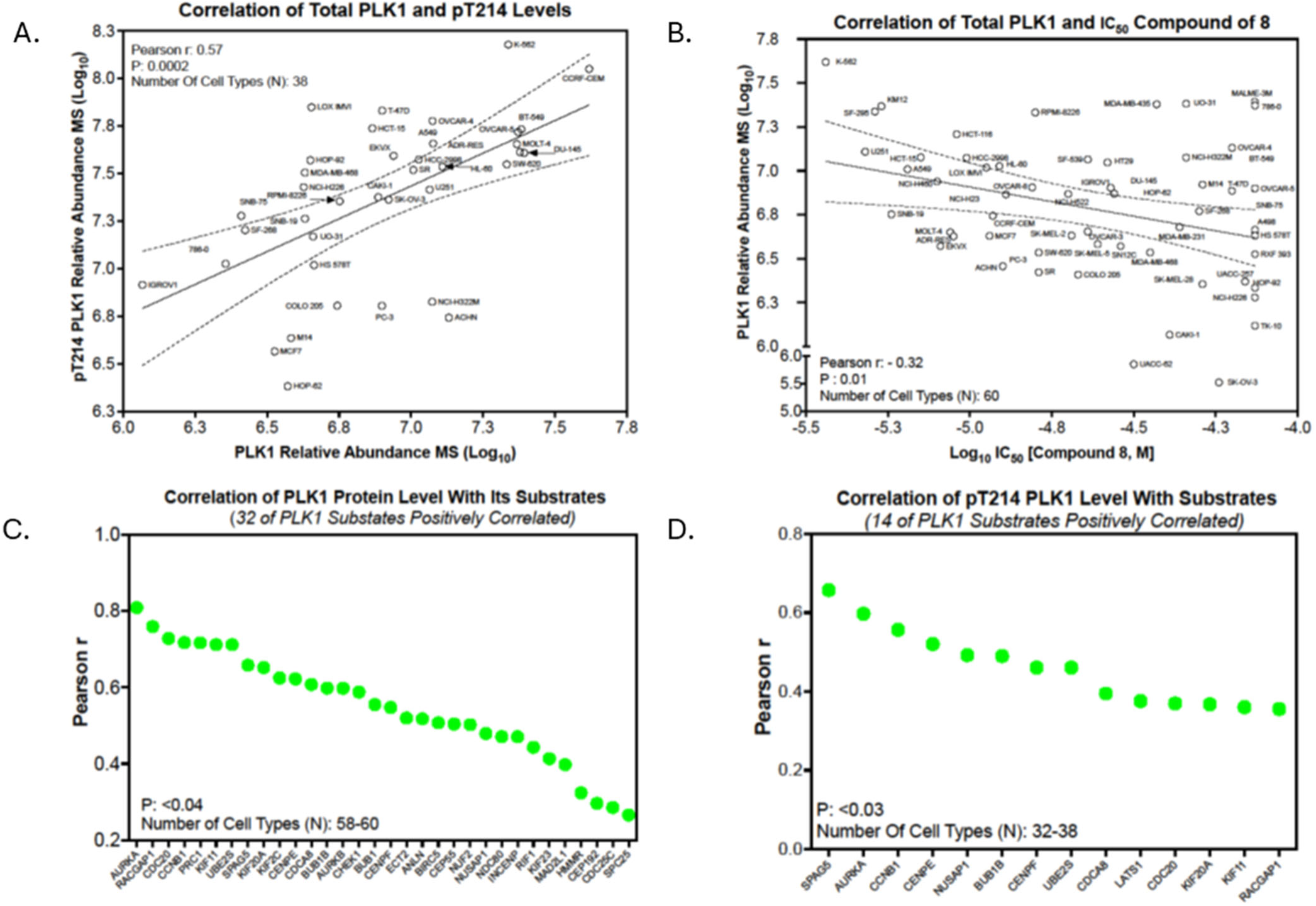
Proteomic analysis of cell lines in the NCI-60 panel. A) Correlation of pT214 PLK1 abundance with total PLK1 abundance across panel of cancer cell lines. B) Correlation of total PLK1 abundance with the IC_50_ of compound 8. C) Correlation of PLK1 protein levels with its substrates; D) Correlation of pT214 PLK1 level with its substrates.

To further explore the biological relevance of PLK1 abundance, correlations were examined between total PLK1 protein levels and known PLK1 substrates from the PhosphoSitePlus database. Total PLK1 protein quantified in all 60 cell lines by mass spectrometry-based proteomics showed significant correlations with 32 of 55 annotated PLK1 substrates (P < 0.05) (**Figure 3C**). The strongest positive correlations were observed for AURKA (r = 0.81), RACGAP1 (r = 0.76), CDC20 (r = 0.73), CCNB1 (r = 0.72), KIF11 (r = 0.71), and UBE2S (r = 0.71), all of which are key regulators of mitotic progression. Similarly, PLK1 pT214 levels showed significant correlations with 14 substrates, consistent with the strong association between total PLK1 abundance and pT214 levels (**Figure 3D**). By comparison, PLK1 pT210 levels exhibited significant correlations with only two substrates, TP53 (r = −0.56) and ECT2 (r = −0.43), both of which were negative. Given the substantially smaller number of cell lines in which pT210 was detected, these limited associations should be interpreted cautiously and likely reflect reduced statistical power.

### Confirmation of cellular engagement of PLK1 by abbapolins

Previous studies established the ability of abbapolins to cause PLK1 degradation at concentrations (Degradation Concentration, DC_50_) that mirrored the anti-proliferative IC_50_ values in PC3 cells [31]. The ability of **6** and **8** to cause loss of PLK1 in PC3 cells in relation to the apparent delayed onset of cell death observed with these two abbapolins was measured by assessing time dependent degradation. Consistent with published data for previous abbapolin analogs, **8** induced PLK1 loss in PC3 cells at a 24 hour time point, albeit at higher concentrations (data not shown).

Interestingly, PLK1 degradation induced by **6** is clearly observable by 48 and 72 h time points (**Figure 4A**), suggesting a delayed onset of action for both loss of PLK1 and antiproliferative activity, while further confirming on-target activity. A dose response was therefore assessed after 72 h abbapolin treatment. Both abbapolins induced a striking decrease in PLK1 at this time point, with DC_50_ values of 3.186 ± 1.057 μM for **6** and 5.005 ± 2.377 μM for **8,** respectively.

**Figure 4:**
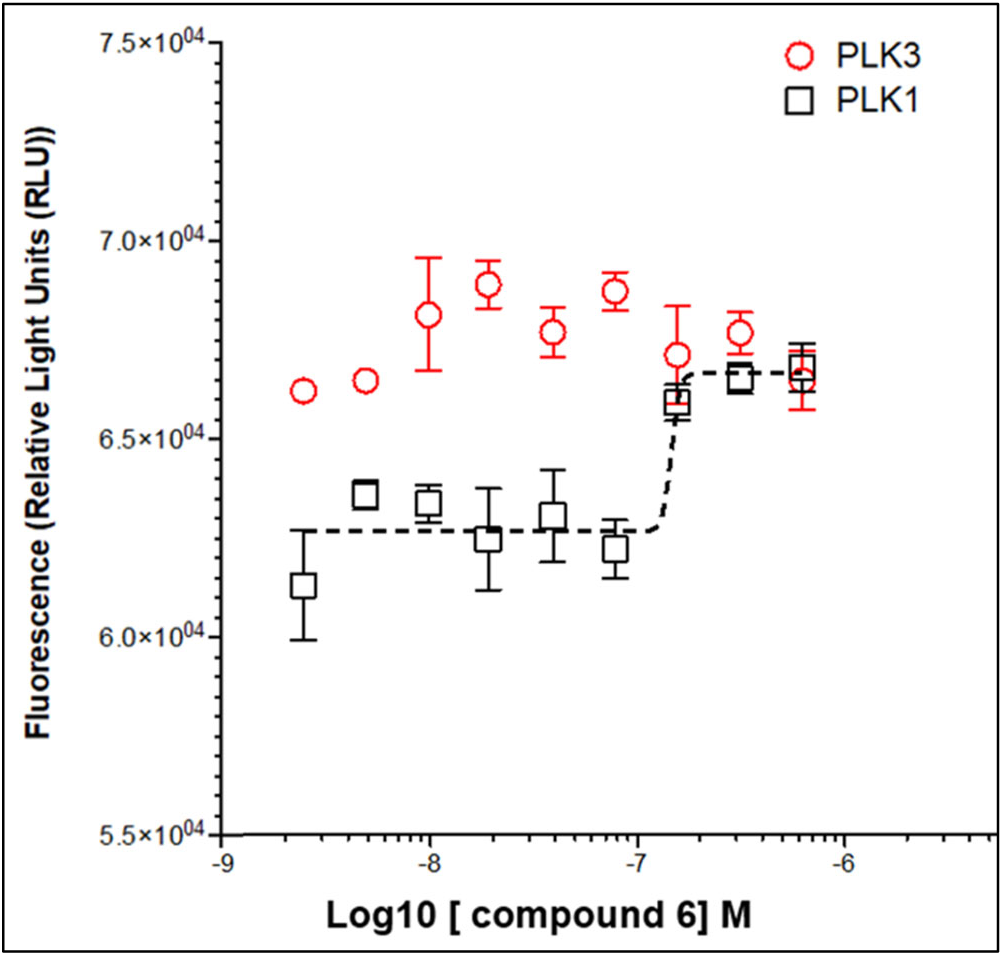
PLK1 degradation in PC3 cells upon treatment with abbapolins. (A). PC3 cells were treated with 30 mM (**6**) for 24, 48, or 72 h. (B) PC3 cells were treated with increasing doses of **6** (left) and **8** (right) for 72 h. (C) Percent of PLK1 detected, normalized to GAPDH. Mean values and standard deviations were obtained from three independent experiments. (D) Plot of dose-dependent PLK1 loss and curve fit to calculate the DC_50_.

Having determined that abbapolin **6** caused a potent loss of cellular PLK1, the direct engagement of cellular PLK1 was then investigated. Using a cellular thermal shift assay (CETSA), it was previously demonstrated that abbapolins show in-cell, on-target PLK1 binding [21, 30, 31]. Here, a novel Micro-Tag^®^ cell target engagement technology (CellarisBio, San Diego) was utilized to determine cellular PLK1 engagement. The data in **Figure 5** show that **6** potently engages PLK1 in cells as determined by an increase in PLK1 thermal stability at sub-micromolar doses, whereas PLK3 stability is unaffected at the same dose range and up to 1 μM, which confirms previous studies with other abbapolins that displayed selectivity for PLK1 over PLK3 [30].

**Figure 5:**
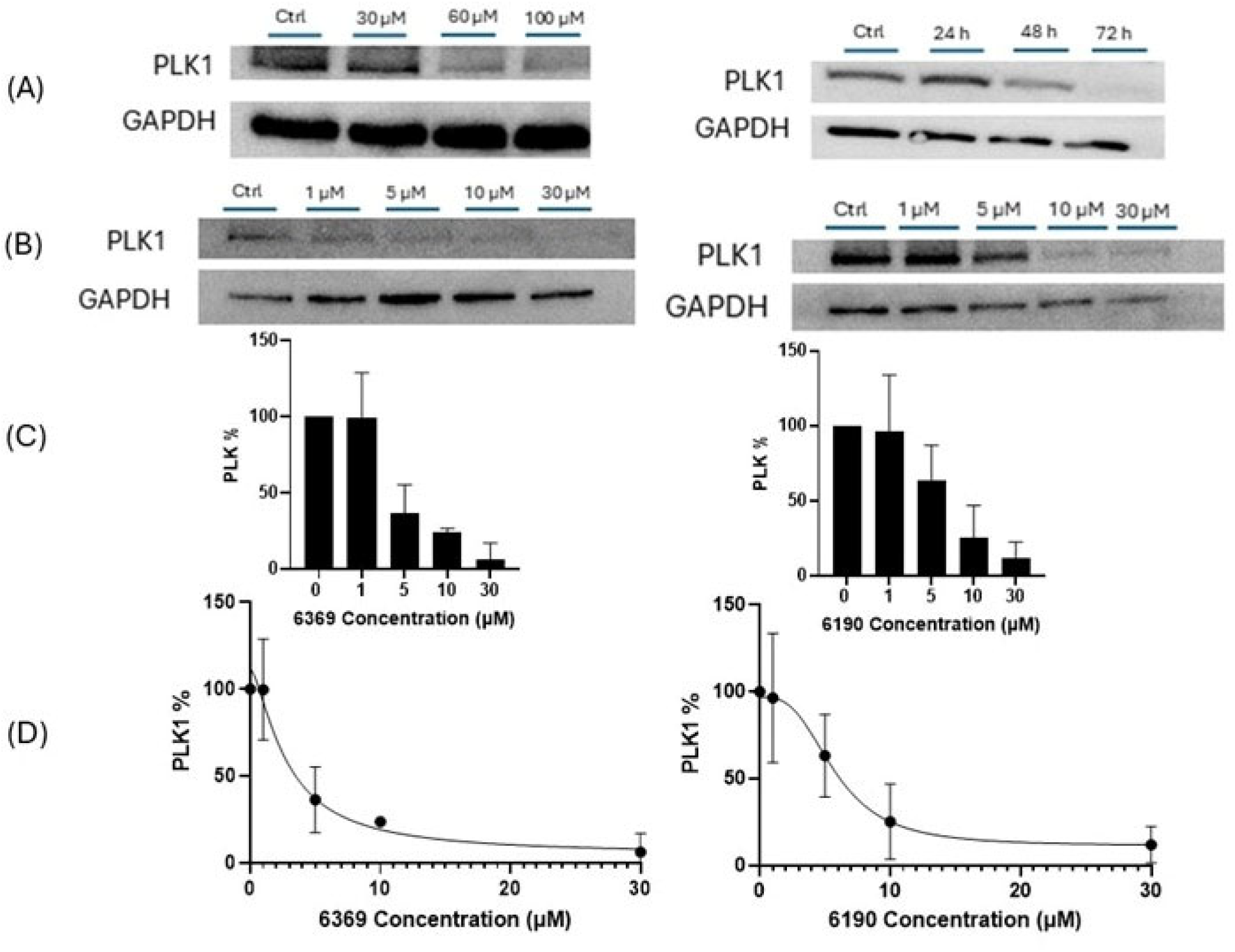
PLK Selectivity of abbapolins: Cellular target engagement of **6** measured by a Micro-Tag^®^ method of PLK thermal stability. Dose dependent stabilization of PLK1 (black squares) is visible at doses above 300 nM. In contrast, no change in stabilization of PLK3 is seen at the same doses.

### In vivo pharmaokinetic analysis and efficacy in protate cancer xenograft

Following the encouraging *in vitro* results, abbapolins **6**, **7**, and **8**, were chosen to evaluate their *in vivo* pharmacokinetic (PK) and anti-tumor efficacy using Swiss Webster mice. The mice were treated at a dose of 50 mg/kg using oral gavage (Table 3). Of these three compounds, **7** was found to have promising PK properties with a C_max_ of 40 μM and a t_1/2_ life of more than 3 h. **8** however had double the C_max_ of **7** and a longer half-life also. **6** was the most potent compound *in vitro* but had approximately 1000- fold lower oral absorption as assessed by C_max_ and AUC (0-8 hrs).

**Table 3.**
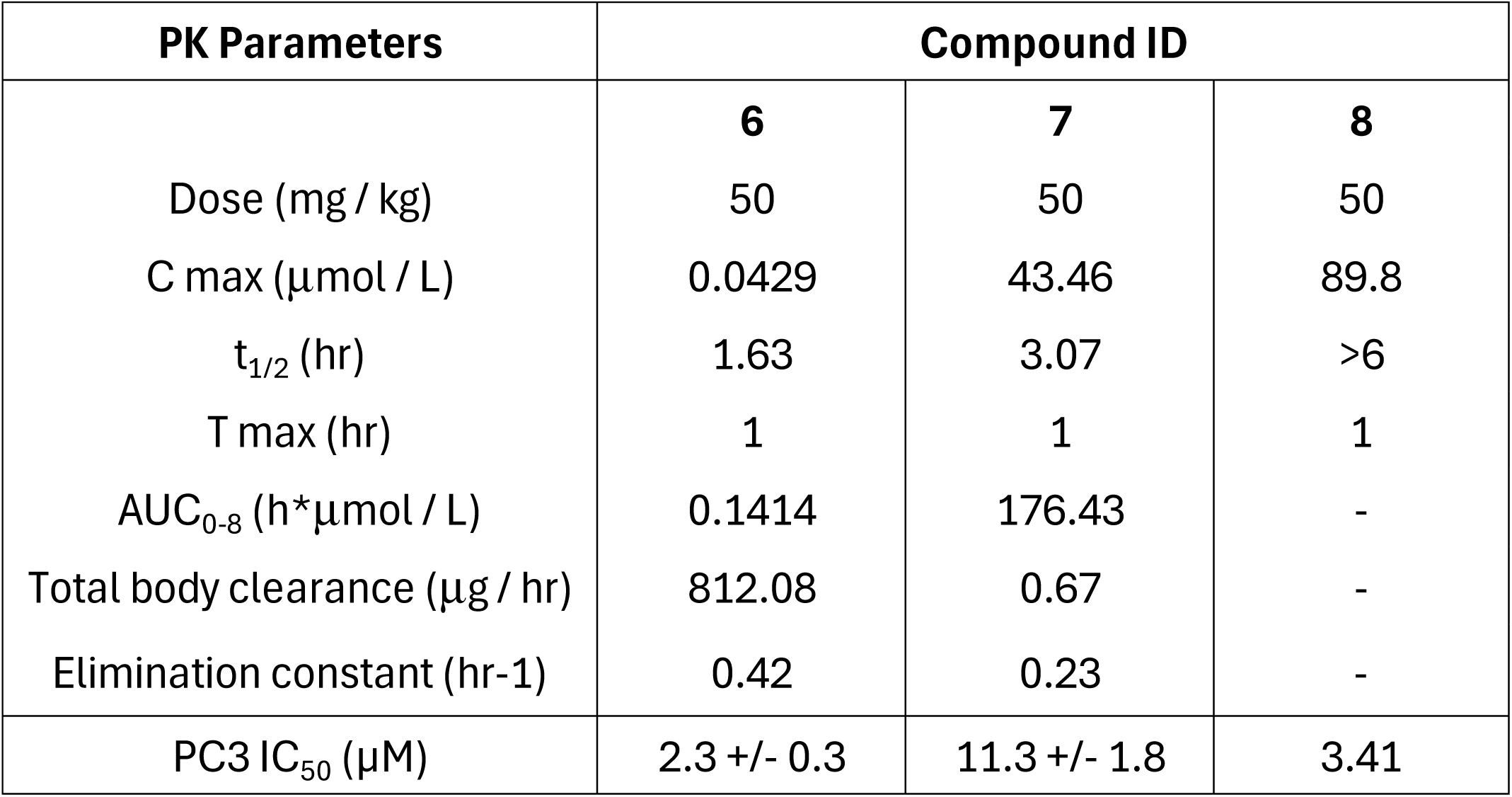
PK properties for orally dosed abbapolins.

| Table 3. PK properties for orally dosed abbapolins |  |  |  |
| --- | --- | --- | --- |
| PK Parameters | Compound ID |  |  |
|  | **6** | **7** | **8** |
| Dose (mg / kg) | 50 | 50 | 50 |
| C max (µmol / L) | 0.0429 | 43.46 | 89.8 |
| t<sub>1/2</sub> (hr) | 1.63 | 3.07 | >6 |
| T max (hr) | 1 | 1 | 1 |
| AUC<sub>0-8</sub> (h*µmol / L) | 0.1414 | 176.43 | - |
| Total body clearance (µg / hr) | 812.08 | 0.67 | - |
| Elimination constant (hr<sup>-1</sup>) | 0.42 | 0.23 | - |
| PC3 IC<sub>50</sub> (µM) | 2.3 +/- 0.3 | 11.3 +/- 1.8 | 3.41 |

A xenograft model of prostate cancer utilizing PC3 cells was employed to determine the *in vivo* activity for **6** and **8**. After establishment of the tumors (200 mm^3^), **6** (55 mg/kg) and **8** (150 mg/kg) were dosed intraperitoneally (i.p.) and orally (p.o.) respectively, once daily for 20 days. The dose of **6** used dictated by its solubility limit in PBS. The *in vivo* efficacy was determined relative to control groups given PBS i.p. or PEG oral dosing vehicles. Statistical analyses revealed no significant differences between the two control groups and, therefore, all untreated mice were combined into a single vehicle control group. As shown in **Figure 6**, tumor volumes were significantly reduced in mice treated with **6** (p= 0.0294) and **8** (p=0.0022) as compared to control mice. Importantly, there was no observable toxicity caused by either abbapolin across the 20-day treatment. The kidneys and livers of all groups were collected and weighed to examine potential adverse effects in these organs. There were no observable weight differences between treatment groups and the vehicle groups for the analyzed organs, which suggests that both abbapolins, **6** and **8** do not cause gross adverse hepatic or nephrotic effects in these treatment conditions and generally are well tolerated by the animals.

**Figure 6.**
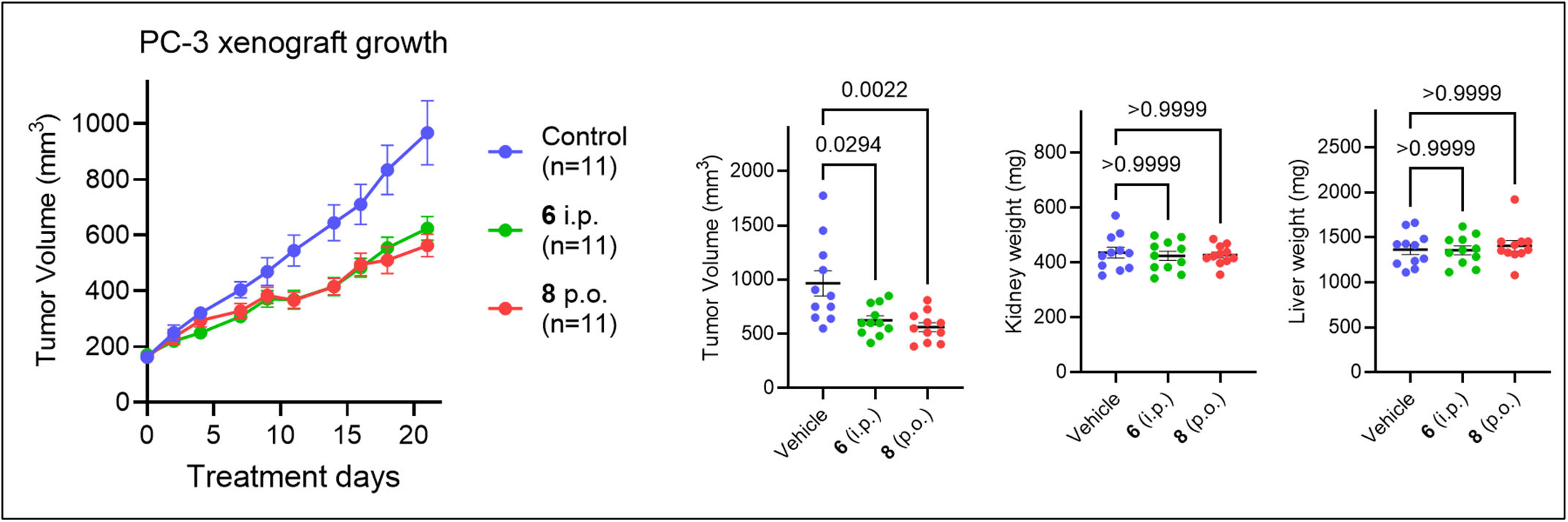
In vivo proof of concept for anti-proliferative activity of abbapolins: PC-3 prostate cancer xenografts carried out in athymic nude mice treated with vehicle control (blue), **6** (green) and **8** (red). The left panel shows tumor volume over the 20-day time course. Next, tumor volume at the end of the experiment is shown for control mice and mice treated with **6** and **8**. The two right panels show organ weights for the kidneys and livers at the end of the experiment for all mice.

An additional xenograft experiment utilizing the 22RV1 prostate cancer cell line was performed using compounds **7** (oral gavage) and **6** (i.p). Oral gavage was chosen for **7** because of its promising pharmacokinetics (100 mg/kg) whereas **6** was again administered i.p. (50 mg/kg) because of its insufficient oral availability in the PK studies (Table 3). Tumor growth inhibition was observed following treatment with **6**, although statistical significance was not reached due to high variability in the untreated control group, whereas **7** did not appear to inhibit tumor growth with this treatment protocol (Supplementary Figure 1). Even though **6** had low oral pharmacokinetic properties compared to **7**, i.p. dosing led to better *in vivo* activity in this model. There was no observable effect on mouse organ or body weight following **6** or **7** treatments, again suggesting tolerance of abbapolins at the doses and treatment times performed (Supplementary Figure 1). The clinically evaluated KDI volasertib was also tested in this xenograft model (50 mg/kg 2 days/week), and it significantly reduced tumor volume (Supplementary Figure 1). However, this ATP competitive inhibitor also caused statistically significant reductions in overall mouse weight as well as spleen, liver, and kidney weights, and one mouse died during the study. These observations are consistent with the toxic effects clinically observed with volasertib. In contrast, no statistically significant changes in body or organ weight were observed with abbapolin treatment in both xenograft experiments.

### Abbapolins lead to loss of PLK1 *in vivo*

Previous studies with abbapolins revealed that these novel PBD-targeting inhibitors induced loss of cellular PLK1, a PROTAC-like degradation phenotype that could be at least partially reversed by proteasome inhibition[31]. To determine if this cellular phenotype was observable *in vivo* and perhaps could serve as a pharmacodynamic marker, the ability of abbapolins to cause loss of PLK1 was measured in tumors (**Figure 7**). Four tumors per group for a total of 16 tumors were analyzed: mice treated with **6**, those treated with PBS vehicle (control for treatment with **6**), mice treated with **8**, and those treated with PEG vehicle (control for treatment with **8**). Tumors were subjected to protein extraction, and lysates were analyzed by Western blotting for PLK1. PLK1 protein levels were shown to be consistently lower in tumors from mice treated with abbapolin compared to the corresponding control tumors (**Figures 7A** and **7B**) and were shown to be statistically significant (**Figure 7C**). Next, the abbapolin-induced decrease in tumor volume (**Figure 6**) was correlated with the decrease in PLK1 protein in these tumors. PLK1 levels normalized to GAPDH were plotted against tumor volume. Tumors from mice treated with **6** and their vehicle treated control (**Figure 7D**) and tumors from compound **8** treated mice and their vehicle treated control (**Figure 7E**) were plotted. Spearman analysis was then conducted to determine the correlation between tumor volume and PLK1 protein for each drug treated group and its vehicle control. A strong correlation was observed for both abbapolins, with a higher correlation between tumor volume and PLK1 protein levels observed with tumors from mice treated with **6** and their control than with tumors from mice treated with **8** and their control.

**Figure 7:**
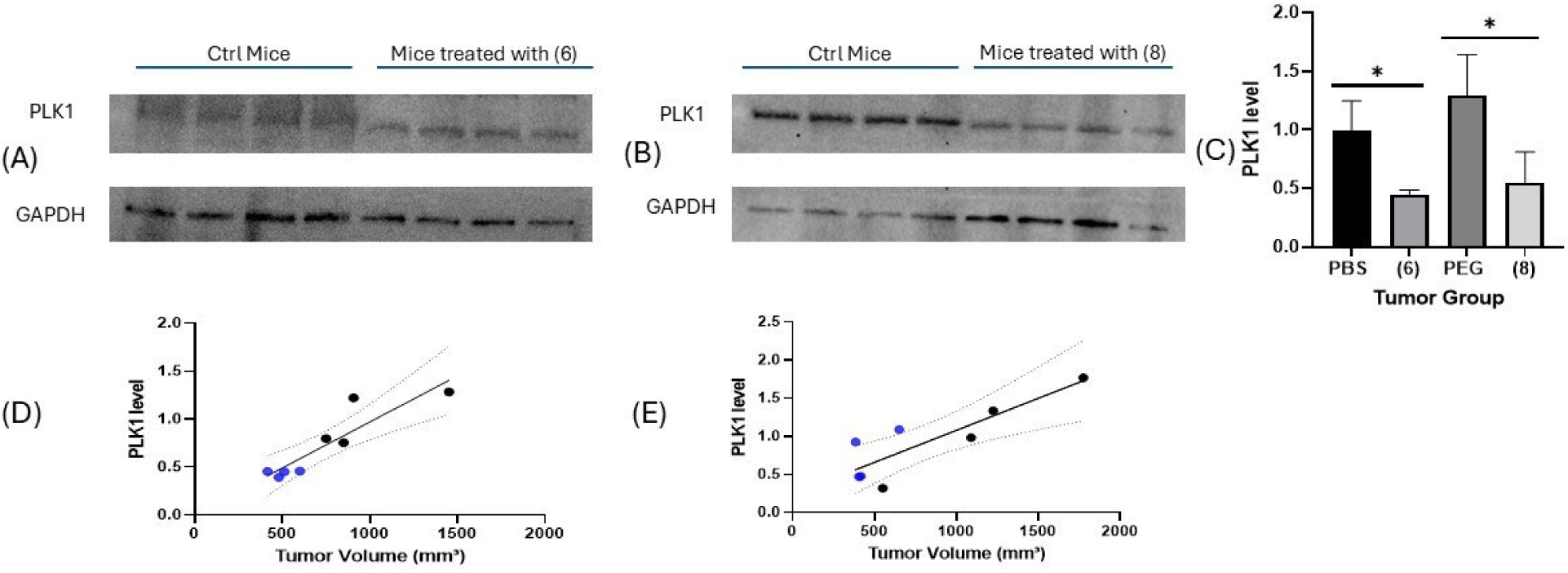
Quantification of PLK1 protein in PC3 prostate cancer xenograft tumors. (A,B) Western Blot images showing PLK1 protein levels relative to GAPDH in tumors from mice treated with **6** or **8** as compared to tumors from mice treated with PBS i.p. (compound **6** control group) or oral PEG (compound **8** control group). (C) Statistical data are presented as mean +/- SD. Three independent experiments were performed and four tumors from each animal group were analyzed. * indicates p<0.05. Unpaired two- sided Welch’s T test was used for statistical analysis. (D) Scatter plot of tumor volume vs. mean PLK1 protein in mice treated with **6** compared to control (in blue) from three independent Western blots per tumor. Line indicates simple linear regression with 95% confidence interval. Correlation was assessed by Spearman rank test: r = 0.9048, p = 0.0046, n = 8 tumors. (E) Scatter plot and analysis as in (D) for mice treated with **8** compared to control: r= 0.7619, p=0.0368, n=8 tumors.

### Enzalutamide resistant tumors are sensitive to abbapolin treatment

The potential of abbapolins as prostate cancer chemotherapeutics was further examined in a different in vitro model, namely androgen resistant prostate cancers. C4-2R cells were treated with the androgen receptor antagonist enzalutamide, abbapolin **6**, or both in combination. As shown in **Figure 8A**, treatment with enzalutamide alone caused minimal change in colony number at a dose of 20 μM, showing that the C4-2R cells are largely unaffected by this dose of anti-androgen treatment. Treatment with **6** alone caused a 50% reduction in colony number at a dose of 20 μM. However, the combination treatment of enzalutamide and **6** at an equimolar 20 μM dose of each produced a strong growth inhibitory effect, nearly eliminating all visible colonies. A heatmap of synergy and 3D synergy landscape were generated to examine the combination of abbapolin and anti-androgen treatment. A score of 15.2 demonstrates strong synergy with the combination and suggests that abbapolins can reverse androgen resistance in prostate cancer.

**Figure 8.**
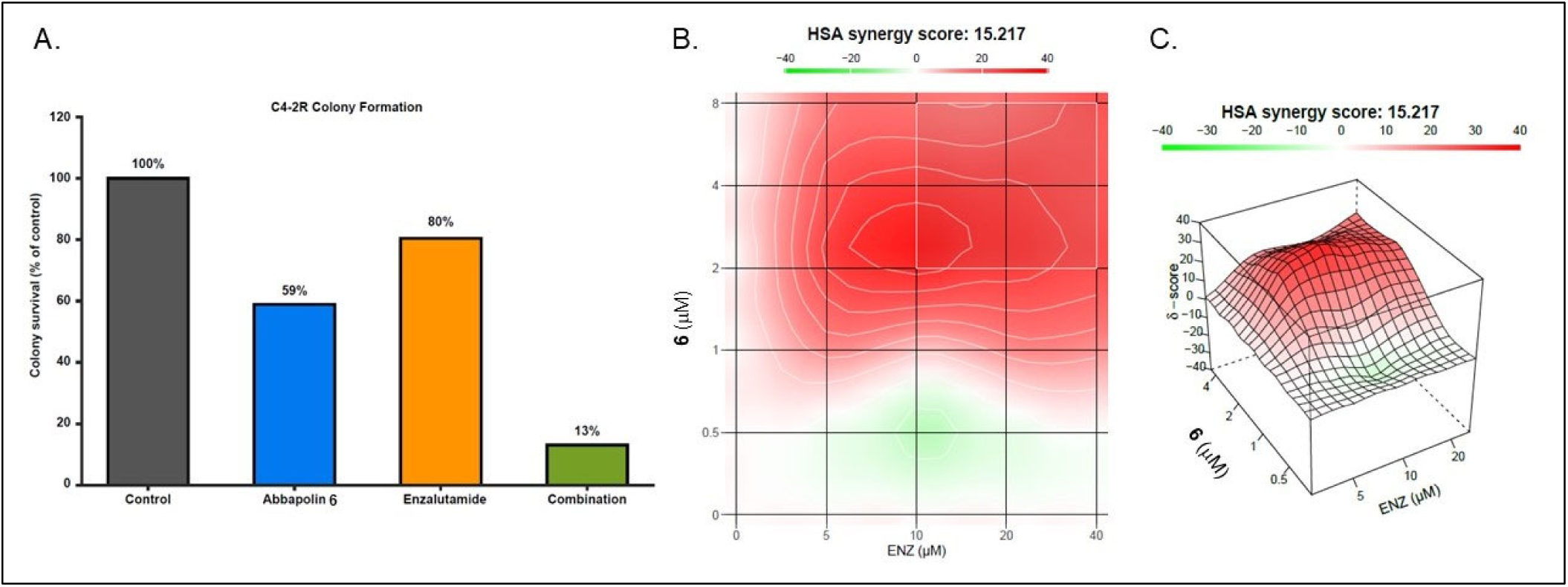
Synergy of abbapolins in combination with enzalutamide. Androgen resistant C4-2R cells were treated with enza alone or with **6**. **8A**, plot of numerical results from colony forming assays with and without enza. **8B**, synergy heatmap with **6** on the y-axis and enza on the x-axis. **8C**, 3D energy landscape plot of synergy with **6** and enza on the x- and y-axes and synergy score on the z-axis.

## DISCUSSION

Polo-like kinase 1 (PLK1) remains one of the most compelling therapeutic targets for the treatment of human cancer because of its central role in regulating mitotic progression and its frequent overexpression in aggressive malignancies. Despite decades of drug development, ATP-competitive PLK1 inhibitors have dose-limiting toxicities, incomplete therapeutic benefit, and the potential emergence of resistance associated with kinase-domain mutations. These limitations have prompted renewed interest in alternative strategies that inhibit PLK1 through mechanisms distinct from catalytic-site blockade. In the present study, we demonstrate that selective targeting of the Polo-box domain (PBD) with optimized abbapolins produces potent antitumor activity in vitro and in vivo while inducing degradation of endogenous PLK1, thereby establishing PBD inhibition as a viable therapeutic strategy for prostate cancer. First generation abbapolins developed using REPLACE, a strategy for targeting protein-protein interactions, provided proof of principle for targeting the PBD [30, 31]. While previously developed compounds were based solely upon mimicking the interactions between the substrate phospho-threonine with a carboxylate isostere, it was recognized that the interaction between phosphorylated substrates and the PBD through ion pairing involving H538 and K540 could be blocked more efficiently through compounds harboring phosphothreonine isosteres, with the goal being to provide greater electrostatic complementarity with the PBD residues. Testing a panel of phosphate isosteres identified that a meta- substituted abbapolin with a phosphate replacing the carboxylate was the most potent compound at engaging PLK1 in a cellular assay and furthermore inhibiting the growth of PC3 prostate cancer cells. The nanomolar activity observed in colony forming assays and cellular engagement of PLK1 confirms the hypothesis that these analogs are excellent bioisosteres.

The results from the NCI-60 panel data revealed tumor cell line sensitivities, notably in the two prostate cancer cell lines, PC3 and DU145, which along with prior work justified the further exploration of abbapolins as potential prostate cancer therapeutics. COMPARE analyses suggest that PBD-targeted abbapolins possess a unique mechanism of action distinct from notable KD-targeted PLK1 inhibitors such as volasertib and across all FDA-approved oncology agents. The finding that no molecules showed significant Pearson coefficients even across the substantial investigational drugs in the database represents impressive circumstantial evidence that abbapolins possess a unique mechanism of action and that targeting the PBD through small molecules is a valid and distinct approach. Previous reports from this laboratory on the results of a genome-wide gene expression study in which the KDI BI2536 showed the expected expression change in mitotic and cell cycle-related genes, while the abbapolins tested showed expression changes in a few overlapping target gene groups but otherwise displayed distinct gene expression changes from those induced by KD inhibiton, thus also providing evidence of distinct mechanism of action[30]. In the COMPARE analysis with abbapolin **8** it was interesting that the highest comparator among FDA approved drugs, albeit with a non-significant correlation coefficient of 0.42, was the COX-2 inhibitor celecoxib, notable because it was reported that COX-2 inhibitors down regulate kinetochore/centromere proteins including PLK1 in prostate cancer cell lines[35]. Further investigation is required to determine the veracity of the interactions of celecoxib with PLK1.

Importantly, NCI-60 screening revealed key structural features of abbapolins required for potent anti-proliferative activity, namely those containing a thioether or amine group connecting the alkyl tail with the benzamido group and which engages the cryptic pocket in the PBD substrate recognition site, showed the highest activity. As the differences in duration of drug exposure could have impacted the results between the different viability assays for the same abbapolins, a colony formation assay was utilized to evaluate the consequences of longer treatment times and the potential delayed onset of action. Compound **6** showed potent inhibition with an IC_50_ of 340 nM in contrast to the results obtained with shorter term viability assays. One possible explanation for increased potency of **6** upon longer exposure might be delayed cellular penetration due to the presence of the phosphate group. Regardless of potential kinetic differences among abbapolins in uptake and/or clearance, the observations of PLK1 engagement by Micro-Tag® assay, as well as the observed loss of cellular PLK1 at longer time points closer to the onset of cell death, confirm in-cell and on-target activity of these latest abbapolins. Proteomic analysis of the 60 tumor cell lines comprising the NCI panel revealed that an inverse correlation exists between the levels of PLK1 protein and sensitivity to the abbapolins further confirming on target mechanism of action. As hypothesized, the overexpression of PLK1 is essential for tumor survival and confers a vulnerability that can be exploited therapeutically.

Having obtained highly potent antiproliferative activity *in vitro*, the most promising abbapolins were investigated in xenograft models of prostate cancer. A preliminary *in vivo* efficacy experiment was carried out using 22RV1 castration-resistant prostate cancer cells with **6** and **7** (Supplementary Figure 1). Tumor growth inhibition was clearly observed following treatment with abbapolin **6**, although it did not reach statistical significance (Supplementary Figure 1). The clinically evaluated KDI volasertib was also tested in this model system, and significantly reduced tumor volume. However, volasertib treatment also caused statistically significant reductions in overall mouse weight as well as spleen, liver, and kidney weights, and one of the mice died during the study. These observations are consistent with the toxic effects clinically observed with volasertib. In contrast, no statistically significant changes in body or organ weight were observed with abbapolin treatment. Further, **6** and **8**, the most active abbapolins in the NCI-60 screen, were evaluated in a PC3 xenograft and both abbapolins caused a statistically significant reduction in tumor volume. During both studies the body weight of the mice and the organ weight were monitored and shown not to vary compared to the untreated control animals, suggesting that the abbapolins were tolerated at the doses used.

The observations that abbapolins cause PLK1 loss in PC3 cells inspired an investigation *in vivo* in tumors and whether PLK1 protein could act as a potential biomarker of *in vivo* activity. Indeed, a significant reduction in PLK1 was observed in groups of tumors from mice treated with **6** or **8**. Furthermore, the analysis showed that PLK1 reduction correlates linearly with tumor volume and therefore is a promising pharmacodynamic readout of tumor responsiveness, although further studies will be required to validate intratumoral PLK1 as a clinical biomarker of drug efficacy.

Prostate cancer resistance to the androgen receptor antagonist enzalutamide has emerged as a major clinical challenge. Encouragingly, studies have shown that PLK1 inhibition with KDIs represses the androgen signaling pathway in prostate cancer [36, 37]. These observations spurred investigation into whether abbapolins could also overcome resistance to enzalutamide in tumors with aberrant androgen receptor expression. Indeed, strong synergy between abbapolin **6** and enzalutamide was observed in the C4-2R cell line, which suggests that PLK1 inhibition via the PBD disrupts androgen-dependent signaling. It was recently reported that PLK1 promotes the Hedgehog signaling pathway and this drives resistance to enzalutamide [38]. Further studies will be carried out to investigate the connection to Hedgehog signaling and thus provide further insights into how abbapolins might be effective in CRPC.

Collectively, this study of abbapolin anti-tumor efficacy provides *in vivo* proof-of-concept for applying the REPLACE strategy for conversion of potent but non-drug-like peptide ligands into efficacious small molecules both in general for protein-protein interactions and specifically for the PBD of PLK1. Lead abbapolin compounds **6** and **8** have been identified and represent excellent starting points for further development as prostate cancer therapeutics. These compounds also provide validation for targeting the PBD as an alternate strategy to the ATP binding site, which has not yet yielded FDA approved drugs. Several PLK1 KDIs have been abandoned during clinical development due to lack of efficacy and toxicity. The results from this study reveal that there are advantages to targeting PLK1 in cancer via non-ATP competitive means. These include potently and selectively targeting PLK1 versus PLK3, a known tumor suppressor (**Figure 5**). Recent observations suggest molecules that bind to the KD produce an open conformation of PLK1 that may alter the function and intracellular partners of PLK1 through increased affinity for its substrates [21, 22]. This laboratory and others have observed an accumulation of PLK1 in cells treated with KD inhibitors, which can easily be interpreted to result from the natural upregulation of PLK1 in mitosis and the mitotic arrest seen with KD inhibitors. However, the form and function of the catalytically inactive but open conformation of PLK1 is poorly understood. KD inhibitors therefore may not block non-catalytic roles of PLK1 and in fact may promote non-enzymatic functions, which might in part explain the lack of success in advancing KD inhibitors beyond clinical trials. In summary, the abbapolins reported here represent a promising and novel means of targeting PLK1 that has significant potential for combination therapy for androgen-resistant prostate cancer.

## AUTHOR CONTRIBUTIONS

The manuscript writing contributions included GM, GR, CNR, MP, MF, DC, MDW and CM. GM, DC and JS designed and carried out cellular experiments including western blotting, antiproliferative assays. MF, YK and XL designed and carried out cellular combination studies in castrate resistant prostate cancer. GR, JS and CNR designed and synthesized all the abbapolin derivatives described within. GM, KH, NJ and MP contributed to the in vivo study of abbapolin molecules. EN and IB developed and performed the Micro-Tag® PLK1 and PLK3 cellular target engagement assay. MC, ZM, CS and SK contributed to PK studies and data analysis. DC contributed to correlation analysis of phosphoproteomic and PLK1 substrates with abbapolin cellular activity. MDW and CM were responsible for the overall project direction including hypothesis generation, design of abbapolin inhibitors, design and implementation of cellular studies, supervision and manuscript writing.

## DATA AVAILABILITY STATEMENT

All materials described in this manuscript, including all relevant raw data, will be freely available to any researcher wishing to use them for non-commercial purposes, without breaching participant confidentiality.

## ACKNOWLEDGEMENTS

We thank Dr Bill Cotham and Dr Perri Pellechia for providing the expertise and resources for mass spectrometry and nuclear magnetic resonance respectively.

## REFERENCES

1. Barr, F.A., H.H. Sillje, and E.A. Nigg, Polo-like kinases and the orchestration of cell division. Nat Rev Mol Cell Biol, 2004. 5(6): p. 429–40.

2. Archambault, V. and D.M. Glover, Polo-like kinases: conservation and divergence in their functions and regulation. Nat Rev Mol Cell Biol, 2009. 10(4): p. 265–75.

3. Wyatt, M.D. and C. McInnes, Insights into the Structural Regulation of Polo-Like Kinase Activity using AlphaFold. Biorxiv, 2024.

4. Chapagai, D., et al., Structural regulation of PLK1 activity: implications for cell cycle function and drug discovery. Cancer Gene Ther, 2025. 32(6): p. 608–621.

5. Zitouni, S., et al., Polo-like kinases: structural variations lead to multiple functions. Nat Rev Mol Cell Biol, 2014. 15(7): p. 433–52.

6. Raab, M., S. Becker, and M. Sanhaji, Targeting polo-like kinase 1: advancements and future directions in anti-cancer drug discovery. Expert Opin Drug Discov, 2024. 19(10): p. 1153–1157.

7. Strebhardt, K., Multifaceted polo-like kinases: drug targets and antitargets for cancer therapy. Nat Rev Drug Discov, 2010. 9(8): p. 643–60.

8. Hanisch, A., et al., Different Plk1 functions show distinct dependencies on Polo-Box domain-mediated targeting. Mol Biol Cell, 2006. 17(1): p. 448–59.

9. Park, J.E., et al., Polo-box domain: a versatile mediator of polo-like kinase function. Cell Mol Life Sci, 2010. 67(12): p. 1957–70.

10. Lowery, D.M., et al., The Polo-box domain: a molecular integrator of mitotic kinase cascades and Polo-like kinase function. Cell Cycle, 2004. 3(2): p. 128–31.

11. de Carmo Avides, M., A. Tavares, and D.M. Glover, Polo kinase and Asp are needed to promote the mitotic organizing activity of centrosomes. Nature Cell Biology, 2001. 3(4): p. 421–424.

12. Lee, K.S., et al., Plk is an M-phase-specific protein kinase and interacts with a kinesin-like protein, CHO1/MKLP-1. Molecular and Cellular Biology, 1995. 15(12): p. 7143–51.

13. Weichert, W., et al., Polo-like kinase I is overexpressed in prostate cancer and linked to higher tumor grades. Prostate, 2004. 60(3): p. 240–245.

14. Ran, Z., et al., Clinicopathological and prognostic implications of polo-like kinase 1 expression in colorectal cancer: A systematic review and meta-analysis. Gene, 2019. 721: p. 144097.

15. Tut, T.G., et al., Upregulated Polo-Like Kinase 1 Expression Correlates with Inferior Survival Outcomes in Rectal Cancer. PLoS One, 2015. 10(6): p. e0129313.

16. Sur, S., et al., A panel of isogenic human cancer cells suggests a therapeutic approach for cancers with inactivated p53. Proc Natl Acad Sci U S A, 2009. 106(10): p. 3964–9.

17. Luo, J., et al., A genome-wide RNAi screen identifies multiple synthetic lethal interactions with the Ras oncogene. Cell, 2009. 137(5): p. 835–48.

18. Liu, X.S., et al., Polo-like kinase 1 facilitates loss of Pten tumor suppressor-induced prostate cancer formation. The Journal of biological chemistry, 2011. 286(41): p. 35795–800.

19. Helmke, C., S. Becker, and K. Strebhardt, The role of Plk3 in oncogenesis. Oncogene, 2016. 35(2): p. 135– 47.

20. Burkard, M.E., A. Santamaria, and P.V. Jallepalli, Enabling and disabling polo-like kinase 1 inhibition through chemical genetics. ACS Chem Biol, 2012. 7(6): p. 978–81.

21. Chapagai, D., et al., Structural Basis for Variations in Polo-like Kinase 1 Conformation and Intracellular Stability Induced by ATP-Competitive and Novel Noncompetitive Abbapolin Inhibitors. ACS Chem Biol, 2023. 18(7): p. 1642–1652.

22. Raab, M., et al., Modulation of the Allosteric Communication between the Polo-Box Domain and the Catalytic Domain in Plk1 by Small Compounds. ACS Chem Biol, 2018. 13(8): p. 1921–1931.

23. Craig, S.N., M.D. Wyatt, and C. McInnes, Current assessment of polo-like kinases as anti-tumor drug targets. Expert Opin Drug Discov, 2014. 9(7): p. 773–89.

24. Stafford, J.M., M.D. Wyatt, and C. McInnes, Inhibitors of the PLK1 polo-box domain: drug design strategies and therapeutic opportunities in cancer. Expert Opin Drug Discov, 2023. 18(1): p. 65–81.

25. Narvaez, A.J., et al., Modulating Protein-Protein Interactions of the Mitotic Polo-like Kinases to Target Mutant KRAS. Cell Chem Biol, 2017. 24(8): p. 1017–1028 e7.

26. Yuan, J., et al., Polo-Box Domain Inhibitor Poloxin Activates the Spindle Assembly Checkpoint and Inhibits Tumor Growth in Vivo. Am J Pathol, 2011. 179: p. 2091–2099.

27. Reindl, W., et al., Inhibition of polo-like kinase 1 by blocking polo-box domain-dependent protein-protein interactions. Chem Biol, 2008. 15(5): p. 459–66.

28. Archambault, V. and K. Normandin, Several inhibitors of the Plk1 Polo-Box Domain turn out to be non- specific protein alkylators. Cell Cycle, 2017. 16(12): p. 1220–1224.

29. Scharow, A., et al., Optimized Plk1 PBD Inhibitors Based on Poloxin Induce Mitotic Arrest and Apoptosis in Tumor Cells. ACS Chem Biol, 2015. 10(11): p. 2570–9.

30. Craig, S.N., et al., Structure-activity and mechanistic studies of non-peptidic inhibitors of the PLK1 polo box domain identified through REPLACE. Eur J Med Chem, 2022. 227: p. 113926.

31. Chapagai, D., et al., Nonpeptidic, Polo-Box Domain-Targeted Inhibitors of PLK1 Block Kinase Activity, Induce Its Degradation and Target-Resistant Cells. J Med Chem, 2021. 64(14): p. 9916–9925.

32. Ianevski, A., A.K. Giri, and T. Aittokallio, SynergyFinder 3.0: an interactive analysis and consensus interpretation of multi-drug synergies across multiple samples. Nucleic Acids Res, 2022. 50(W1): p. W739– W743.

33. Lasek, A.L., et al., The Functional Significance of Posttranslational Modifications on Polo-Like Kinase 1 Revealed by Chemical Genetic Complementation. PLoS One, 2016. 11(2): p. e0150225.

34. Frejno, M., et al., Proteome activity landscapes of tumor cell lines determine drug responses. Nat Commun, 2020. 11(1): p. 3639.

35. Bieniek, J., et al., COX-2 inhibitors arrest prostate cancer cell cycle progression by down-regulation of kinetochore/centromere proteins. Prostate, 2014. 74(10): p. 999–1011.

36. Zhang, Z., et al., Inhibition of Plk1 represses androgen signaling pathway in castration-resistant prostate cancer. Cell Cycle, 2015. 14(13): p. 2142–8.

37. Zhang, Z., et al., Plk1 inhibition enhances the efficacy of androgen signaling blockade in castration- resistant prostate cancer. Cancer Res, 2014. 74(22): p. 6635–47.

38. Zhang, Q., et al., The kinase PLK1 promotes Hedgehog signaling-dependent resistance to the antiandrogen enzalutamide in metastatic prostate cancer. Sci Signal, 2025. 18(878): p. eadi5174.

