## Supplemental methods and figures for "Abbapolin inhibitors of the PLK1 PBD as Prostate Cancer Therapeutics, in vivo activity and synergy with androgen therapy"

**Experimental methods and analytical data.**

**Synthesis of 2-(4-octylbenzamido)phenyl dihydrogen phosphate, 1.**

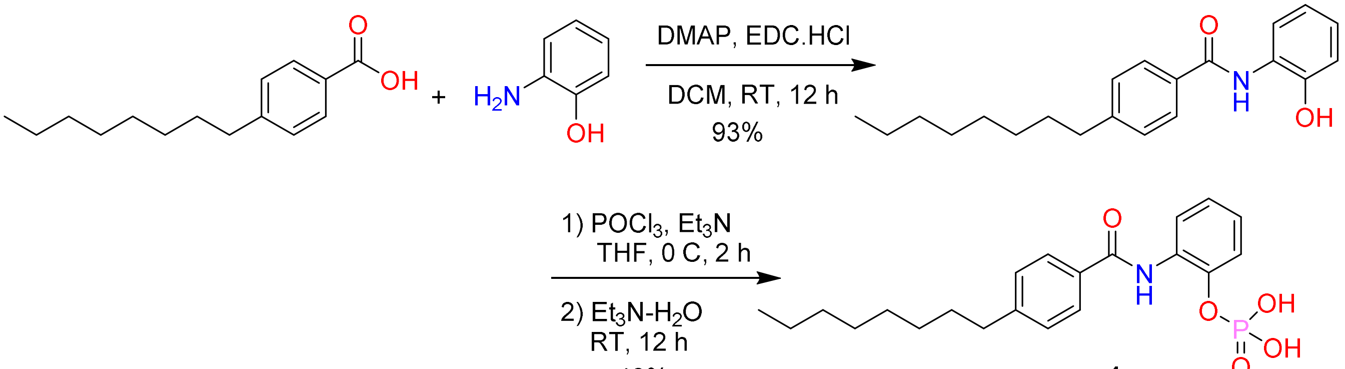


**[2-[(4-octylbenzoyl)amino]phenyl]dihydrogen phosphate (1)**: To a cooled solution of $POCl_3$ (51 μL, 0.553 mmol) in THF (1 ml), a solution of *N*-(2-hydroxyphenyl)-4-octylbenzamide (60 mg, 0.184 mmol) and $Et_3N$ (51 μL, 0.369 mmol) in THF (2 ml) was added slowly over 10 min. The mixture was stirred at 0-5 °C for additional 2 h and warmed to room temperature for additional 1 h. The mixture was slowly quenched by 1:1 water-$Et_3N$ (0.25 ml/0.25 ml) in THF (1 ml) maintaining 0-5 °C. Upon complete quenching, the mixture was concentrated under reduced pressure. The crude was purified by reverse phase C18 combi flash chromatography using 5–95% MeCN-water gradient. The fractions containing the product were pooled and concentrated under reduced pressure to remove MeCN and lyophilized to give [2-[(4-octylbenzoyl)amino]phenyl] dihydrogen phosphate (30 mg, 40%) as a white solid. $^1$H-NMR (300 MHz, DMSO-d6) δ 10.96 (s, 1H), 7.99 (d, *J* = 8.0 Hz, 2H), 7.74 (d, *J* = 7.9 Hz, 1H), 7.33 (d, *J* = 8.0 Hz, 2H), 7.26 (dd, *J* = 9.5, 7.3 Hz, 2H), 7.12 (t, *J* = 7.4 Hz, 1H), 2.65 (t, *J* = 7.6 Hz, 2H), 1.60 (s, 2H), 1.38-1.14 (m, 10H), 0.94-0.76 (m, 3H). $^{31}$P-NMR (121 MHz, DMSO-D6) δ 22.90. ESI-MS (pos): *m/z* 406 $(M+H)^+$.

***N*-(2-Hydroxyphenyl)-4-octyl-benzamide**: To a solution of 4-octylbenzoic acid (200 mg, 0.853 mmol), EDCI-HCl (245 mg, 1.28 mmol) and DMAP (10.4 mg, 0.085 mol) in DCM (2 mL), 2-aminophenol (140 mg, 1.28 mol) in DCM (1 ml) was slowly added at room temperature and the reaction mixture was stirred at room temperature for 16 h. Upon completion of the reaction confirmed by TLC, the reaction mixture was diluted with DCM (5 ml) and washed with 0.5 M HCl (3 ml), sat. $NaHCO_3$ (3 ml), water (3 ml) and brine solution (3 ml). The organic phase was dried over anhydrous $Na_2SO_4$, and solvent was removed under reduced pressure. The crude was purified by $SiO_2$ flash column chromatography using 0-50% ethyl acetate in hexanes to give *N*-(2-Hydroxyphenyl)-4-octylbenzamide (260 mg, 93%) as a white solid. ESI-MS (pos): *m/z* 326 $(M+H)^+$.

**Synthesis of 5-methyl-2-(4-octylbenzamido)phenyl dihydrogen phosphate, 2.**

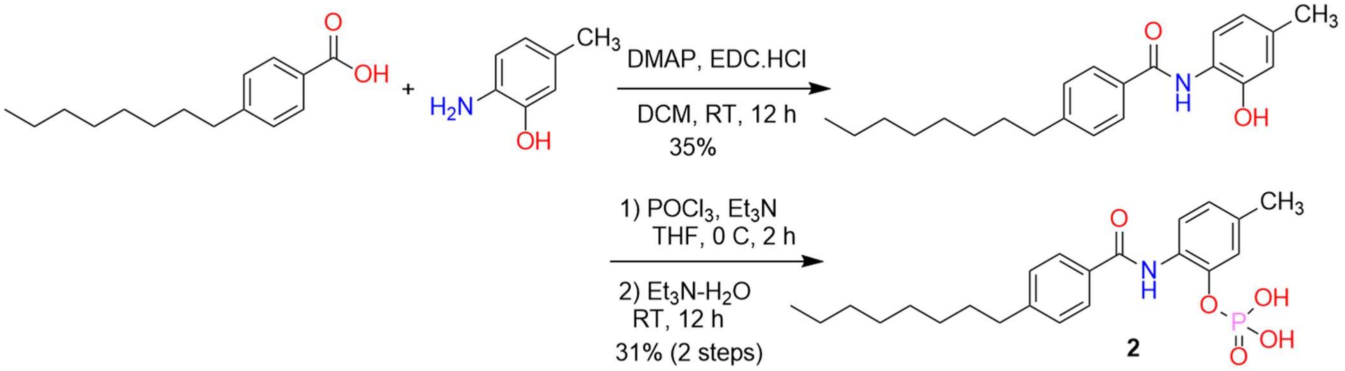


**[5-Methyl-2-[(4-octylbenzoyl)amino]phenyl]dihydrogen phosphate (2)**: To a cooled solution of $POCl_3$ (65 μL, 0.707 mmol) in THF (1 ml), a solution of *N*-(2-hydroxy-4-methylphenyl)-4-octylbenzamide (80 mg, 0.236 mmol) and $Et_3N$ (0.13 ml, 0.943 mmol) in THF (3 ml) was added slowly over 10 min. The mixture was stirred at 0-5 °C for additional 2 h and warmed to room temperature for additional 1 h. The mixture was slowly quenched by 1:1 water-$Et_3N$ (0.25 ml/0.25 ml) in THF (1 ml) maintaining 0-5 °C. Upon complete quenching, the mixture was concentrated under reduced pressure. The crude was purified by reverse phase C18 combiflash chromatography using 5-95% MeCN-water gradient. The fractions containing the product were pooled and concentrated under reduced pressure to remove MeCN and lyophilized to give [5-methyl-2([4-octylbenzoyl]amino]phenyl]dihydrogen phosphate (98 mg, 31%) as a white solid. $^1$H-

NMR (300 MHz, $CDCl_3$) δ 10.07 (s, 1H), 8.02-7.96 (m, 2H), 7.80 (d, *J* = 8.1 Hz, 1H), 7.31-7.21 (m, 3H), 7.15-7.02 (m, 2H), 3.11 (d, *J* = 21.0 Hz, 2H), 2.63-2.54 (m, 2H), 1.56 (t, *J* = 7.5 Hz, 2H), 1.20 (dd, *J* = 11.2, 7.7 Hz, 13H), 0.84-0.78 (m, 3H). $^{31}$P-NMR (121 MHz, $CDCl_3$) δ 28.00. ESI-MS (pos): *m/z* 420 $(M+H)^+$.

***N*-(2-hydroxy-4-methyl-phenyl)-4-octyl-benzamide**: To a solution of 4-octylbenzoic acid (170 mg, 0.725 mmol), EDCI-HCl (209 mg, 1.09 mmol) and DMAP (44 mg, 0.363 mol) in DCM (2 mL), 2-amino-5-methylphenol (134 mg, 1.09 mol) in DCM (1 ml) was slowly added at room temperature and the reaction mixture was stirred at RT for 16 h. Upon reaction completion confirmed by TLC, the mixture was diluted with DCM (5 ml) and washed with 0.5 M HCl (3 ml), sat. $NaHCO_3$ (3 ml), water (3 ml) and brine solution (3 ml). The organic phase was dried over anhydrous $Na_2SO_4$ and solvent was removed under reduced pressure. The crude was purified by $SiO_2$ flash column chromatography using 0–50% EtOAc-hexanes to give *N*-(2-hydroxy-4-methylphenyl)-4-octylbenzamide (85 mg, 35%) as a white solid. ESI-MS (pos): *m/z* 340 $(M+H)^+$.

**Synthesis of (2-(4-octylbenzamido)benzyl)phosphonic acid, 3.**

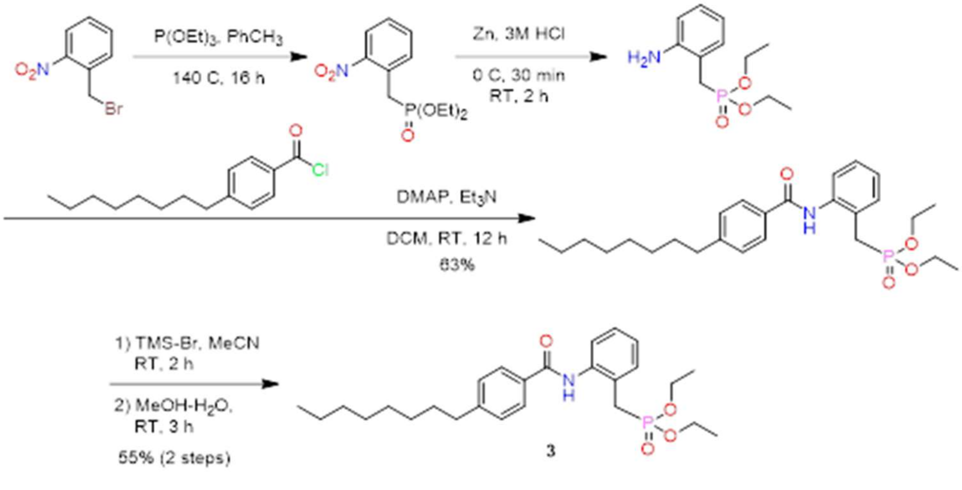


**[2-[(4-Octylbenzoyl)amino]phenyl]methylphosphonic acid, 3**. TMS-Br (0.2 ml, 1.52 mmol) was added to a solution of *N*-[2-(diethoxyphosphorylmethyl)phenyl]-4-octylbenzamide (70 mg, 0.152 mmol) in MeCN (5 ml) at 0 °C. The mixture was warmed to room temperature and stirred for 2 h and concentrated. The residue was slowly quenched with 90% $CH_3OH/H_2O$ (10 ml) at room temperature for 3 h and concentrated under reduced pressure. The crude was purified by reverse phase C18 combiflash chromatography using 5–95% MeCN-water gradient. The fractions containing the product were pooled and concentrated under reduced pressure to remove MeCN and lyophilized to give [2-[(4-octylbenzoyl)amino]phenyl]methyl-phosphonic acid (34 mg, 55%) as a white solid. 1H-NMR (300 MHz, DMSO-D6) δ 10.96 (s, 1H), 7.99 (d, *J* = 8.0 Hz, 2H), 7.74 (d, *J* = 7.9 Hz, 1H), 7.33 (d, *J* = 8.0 Hz, 2H), 7.26 (dd, *J* = 9.5, 7.3 Hz, 2H), 7.12 (t, *J* = 7.4 Hz, 1H), 3.04 (d, *J* = 20.8 Hz, 2H), 2.65 (t, *J* = 7.6 Hz, 2H), 1.59 (s, 2H), 1.35-1.17 (m, 10H), 0.93-0.76 (m, 3H). $^{31}$P-NMR (121 MHz, $CDCl_3$) δ 22.90. ESI-MS (pos): *m/z* 404 $(M+H)^+$.

***N*-[2-(diethoxyphosphorylmethyl)phenyl]-4-octylbenzamide**. To a solution of (diethoxyphosphorylmethyl)aniline (60 mg, 0.247 mmol), DMAP (30 mg, 0.247 mmol) in DCM (4 mL) was added $Et_3N$ (70 µL, 0.493 mmol) followed by the addition of 4-octylbenzoyl chloride (93.5 mg, 0.37 mmol) in DCM (1 mL) and the reaction mixture was stirred at room temperature for 12 h. The reaction was diluted with DCM, washed with 3-M HCl, sat. $NaHCO_3$, water, brine and dried over anhydrous $Na_2SO_4$. DCM was removed under reduced pressure, and the crude was purified by $SiO_2$ flash chromatography (0–50% EtOAc-hexanes) to afford methyl *N*-[2-(diethoxyphosphorylmethyl)phenyl]-4-octylbenzamide (72 mg, 63%) as a yellow solid. ESI-MS (pos): *m/z* 460 $(M+H)^+$.

**Synthesis of (2-(4-octylbenzamido)phenyl)phosphonic acid, 4.**

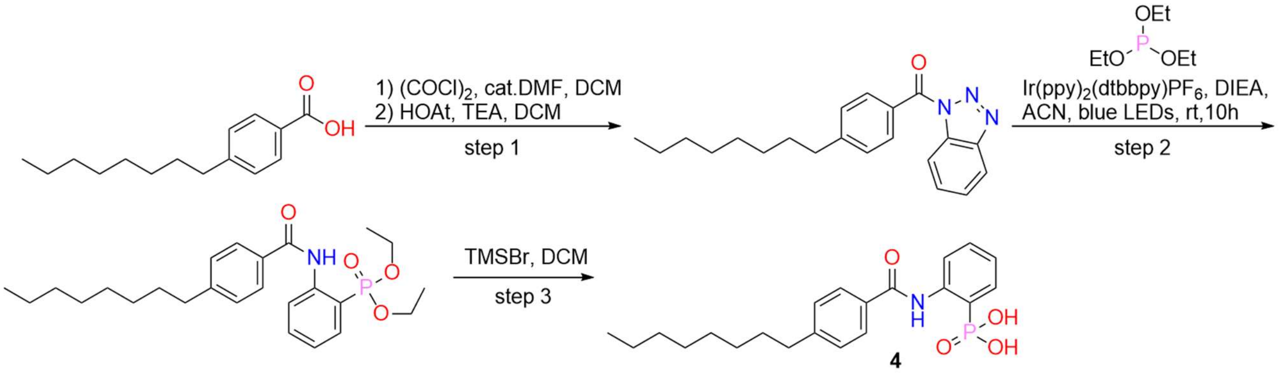


**2-(4-octylbenzamido)phenylphosphonic acid, 4.** To a stirred mixture of diethyl 2-(4-octylbenzamido)phenylphosphonate (100 mg, 0.22 mmol, 1.00 equiv) in DCM (1.00 mL) was added bromotrimethylsilane (343 mg, 2.24 mmol, 10.0 equiv). The mixture was stirred for 20 h at 25 °C. The mixture was concentrated under reduced pressure, the residue was re-dissolved in MeOH/$H_2O$ (9/1, 5 mL) and the resulting mixture was stirred for additional 1 h. The mixture was then concentrated under reduced pressure, the residue was purified by prep-HPLC with the following condition: column: XBridge Shield RP18 OBD column, 19 x 250 mm, 10 um; mobile phase A: water (0.05% TFA ), mobile phase B: ACN; flow rate: 25 mL/min; gradient: 65 B to 95 B in 7 min; 220 nm; RT: 5.22. The collected fraction was partially concentrated under reduced pressure to remove acetonitrile, the precipitated solids were collected by filtration, washed with water (2 x 20 mL) and dried under IR lamp to afford 2-(4-octylbenzamido)phenylphosphonic acid (14.3 mg, 16%) as a white solid. $^1$H-NMR (300 MHz DMSO-$d_6$) δ 12.10 (s, 1H), 11.89 (br s, 1H), 8.68-8.64 (m, 1H), 7.95 (d, *J* = 8.4 Hz, 2H), 7.69-7.62 (m, 1H), 7.58-7.52 (m, 1H), 7.38 (d, *J* = 8.4 Hz, 2H), 7.21-7.16 (m, 1H), 2.69-2.63 (m, 2H), 1.64-1.56 (m, 2H), 1.30-1.21 (m, 10 H), 0.88-0.84 (m, 3H). LCMS (ES, *m/z*): 390 $[M+H]^+$.

**1-(4-octylbenzoyl)-1,2,3-benzotriazole.** To a stirred mixture of 4-octylbenzoic acid (200 mg, 0.85 mmol, 1.00 equiv.) and cat. DMF (1 drop) in DCM (10.0 mL) was added oxalyl dichloride (2.17 g, 17.1 mmol, 20.0 equiv.) dropwise at 0 °C. The mixture was stirred for 5 h at 25 °C and then concentrated under reduced pressure to afford the fresh prepared acyl chloride. The fresh prepared acid chloride was dissolved in DCM (2 mL) and then dropwise into a stirred mixture of benzotriazole (101 mg, 0.85 mmol, 1.00 equiv.) and TEA (345 mg, 3.41 mmol, 4.00 equiv.) in DCM (10.0 mL) at 0 °C. The resulting mixture was stirred for 16 h at 25 °C. The mixture was diluted with DCM (20 mL), washed with water (20 mL) and brine (20 mL), dried over anhydrous sodium sulfate and concentrated under reduced pressure. The residue was purified by silica gel column chromatography, eluted with PE/EtOAc (4:1) to afford 1-(4-octylbenzoyl)-1,2,3-benzotriazole (175 mg, 58%) as a colorless oil. LCMS (ES, *m/z*): 336 $[M+H]^+$

**Diethyl 2-(4-octylbenzamido)phenylphosphonate.** A mixture of 1-(4-octylbenzoyl)-1,2,3-benzotriazole (175 mg, 0.52 mmol, 1.00 equiv), triethyl phosphite (260 mg, 1.56 mmol, 3.00 equiv), DIEA (202 mg, 1.56 mmol, 3.00 equiv) and $[Ir(dtbbpy)(ppy)_2][PF_6]$ (24 mg, 0.03 mmol, 0.05 equiv) was stirred for 10 h at 25 °C under the irradiation of blue LEDs (365 nm). The mixture was concentrated under reduced pressure. The residue was purified by silica gel column chromatography, eluted with PE/EtOAc (1:1) to afford diethyl 2-(4-octylbenzamido)phenylphosphonate (100 mg, 40%) as a colorless oil. LCMS (ES, *m/z*): 446 $[M+H]^+$

**Synthesis of 3-(4-octylbenzamido)phenylphosphonic acid, 5.**

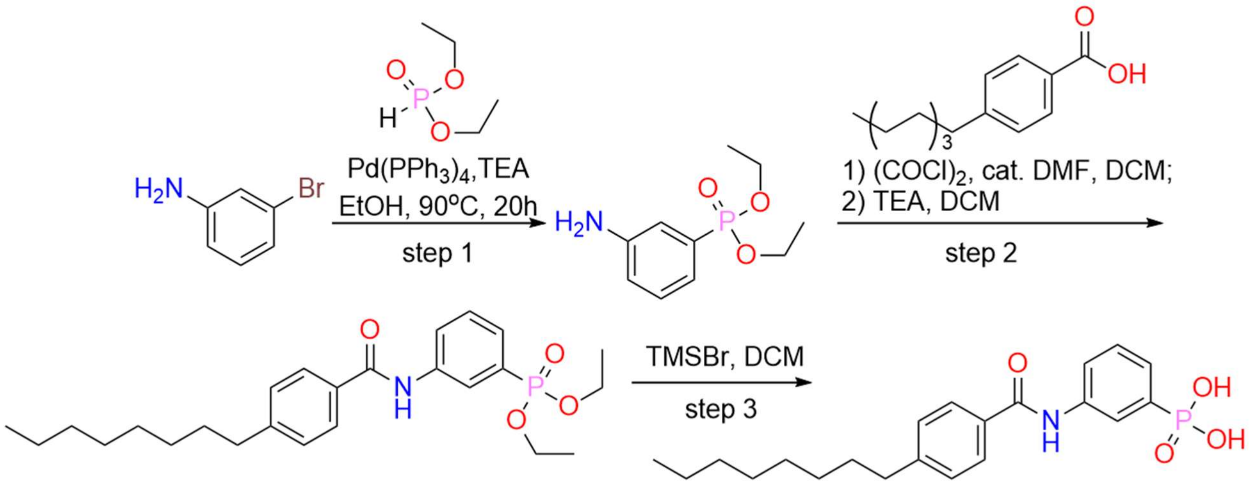


**5**

**3-(4-octylbenzamido)phenylphosphonic acid, 5.** To a stirred mixture of diethyl 3-(4-octylbenzamido)phenylphosphonate (100 mg, 0.22 mmol, 1.00 equiv) in DCM (1.00 mL) was added bromotrimethylsilane (343 mg, 2.24 mmol, 10.0 equiv). The resulting mixture was stirred for 20 h at 25 °C. The mixture was concentrated under reduced pressure, the residue was re-dissolved in MeOH/$H_2O$ (9/1, 5 mL) and the resulting mixture was stirred for additional 1 h. The mixture was then concentrated under reduced pressure, the solids were washed with water (3 x 10 mL) and dried under IR lamp to afford 3-(4-octylbenzamido)phenylphosphonic acid (13.4 mg, 15%) as an off-white solid. $^1$H-NMR (300 MHz DMSO-$d_6$) δ 11.15 (br s, 1H), 10.30 (s, 1H), 8.16 (d, *J* = 13.8 Hz, 1H), 7.96-7.89 (m, 3H), 7.47-7.34 (m, 4H), 2.68-2.63 (m, 2H), 1.62-1.58 (m, 2H), 1.29-1.21 (m, 10H), 0.88-0.79 (m, 3H). LCMS (ES, *m/z*): 390 $[M+H]^+$.

**Diethyl 3-aminophenylphosphonate.** A mixture of m-bromoaniline (300 mg, 1.74 mmol, 1.00 equiv), diethyl phosphonate (289 mg, 2.09 mmol, 1.20 equiv), TEA (352 mg, 3.49 mmol, 2.00 equiv) and $Pd(PPh_3)_4$ (201 mg, 0.17 mmol, 0.10 equiv) in EtOH (6 mL) was stirred for 20 h at 90 °C under nitrogen atmosphere. The mixture was cooled to room temperature and concentrated under reduced pressure. The residue was purified by silica gel column chromatography, eluted with PE/EtOAc (1:9) to afford diethyl 3-aminophenylphosphonate (150 mg, 35%) as a yellow oil. LCMS (ES, m/z): 230 $[M+H]^+$

**Diethyl 3-(4-octylbenzamido)phenylphosphonate.** To a stirred mixture of 4-octylbenzoic acid (153 mg, 0.65 mmol, 1.00 equiv) in DCM (6.00 mL) was added cat. DMF (1 drop) and oxalyl dichloride (1661 mg, 13.1 mmol, 20.0 equiv) dropwise at 0 °C. The mixture was stirred for 5 h at 25 °C and then concentrated under reduced pressure. The fresh prepared acyl chloride was re-dissolved in DCM (2 mL) and then dropwise into a stirred mixture of diethyl 3-aminophenylphosphonate (150 mg, 0.65 mmol, 1.00 equiv) and TEA

(265 mg, 2.62 mmol, 4.00 equiv) in DCM (6.00 mL) at 0 °C. The resulting mixture was stirred for 16 h at 25 °C. The mixture was diluted with DCM (20 mL), washed with water (20 mL) and brine (20 mL), dried over anhydrous sodium sulfate and concentrated under reduced pressure. The residue was purified by silica gel column

chromatography, eluted with PE/EtOAc (4:1) to afford diethyl 3-(4

octylbenzamido)phenylphosphonate (100 mg, 31%) as a yellow solid. LCMS (ES, m/z): 446 $[M+H]^+$

**Synthesis of 2-methyl-5-(4-(octyloxy)benzamido)phenyl dihydrogen phosphate (6).**

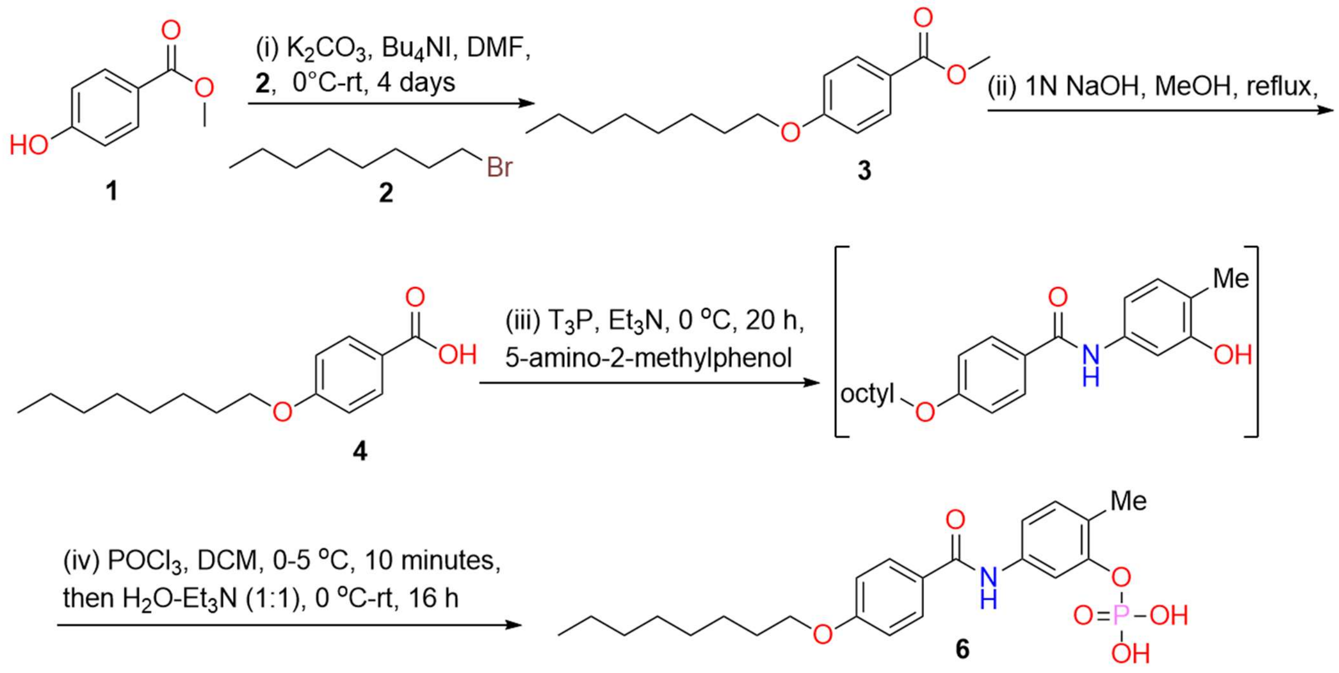


$T_3P$ (2.54 g, 7.99 mmol, 50% wt. in EtOAc) is added to a mixture of 4-Octyloxybenzoic acid (1.00 g, 3.99 mmol) and 5-Amino-O-cresol (0.59 g, 4.79 mmol) in TEA (1.11 mL, 7.99 mmol) and DCM, and the resulting homogeneous solution was stirred at 0 °C for around 20 hours. During the reaction the temperature was slowly brought to rt (24 °C). After 20 hours, reaction was quenched with 3N HCl. A white precipitate was formed immediately. The precipitation was filtered and washed with water several times to remove undesired biproducts. The collected white solids (1.5 g) were dried under vacuum and used for the subsequent step without further purification.

To a cooled solution of $POCl_3$ (1.2 mL, 13.1 mmol) in THF (1 ml) a solution of N-(3-hydroxy-4-methyl-phenyl)-4-octoxy-benzamide intermediate (1.5 g, 4.22 mmol) and TEA (2.46 mL, 17.6 mmol) in THF (3 mL) was added slowly over 10 min. The mixture was stirred at 0-5 °C for additional 2 h and warmed to room temperature for additional 1 h. The mixture was slowly quenched by 1:1 water-$Et_3N$ (0.5 mL) in THF (1 mL) maintaining 0-5 °C. Ice melted overnight. Reaction was monitored by LCMS. Concentrated the reaction mixture and purified the crude material by MPLC reverse-phase chromatography using C-18 column with 1% aqueous formic acid and acetonitrile system as mobile phase. Concentrated fractions gave a slimy thick liquid which was lyophilized to give a fine white powder of **6** (0.75 g, yield 43%) in its pure form. $^{1}$H-NMR (400 MHz, DMSO) δ 10.10 (s, 1H), 7.94 (d, *J* = 8.6 Hz, 2H), 7.74 (s, 1H), 7.48 (dd, *J* = 8.3, 2.2 Hz, 1H), 7.14 (d, *J* = 8.3 Hz, 1H), 7.02 (d, *J* = 8.7 Hz, 2H), 4.04 (t, *J* = 6.5 Hz, 2H), 2.19 (s, 3H), 1.80-1.67 (m, 2H), 1.42 (dd, *J* = 9.6, 5.7 Hz, 2H), 1.37-1.20 (m, 8H), 0.93-0.76 (m, 3H). $^{13}$C-NMR (101 MHz, DMSO) δ 165.2, 161.8, 150.3, 138.4, 130.7, 130.0, 127.2, 124.6, 116.5, 114.4, 113.3, 68.2, 31.7, 29.2, 29.1, 29.1, 26.0, 22.6, 16.3, 14.4. $^{31}$P-NMR (162 MHz, DMSO) δ -5.95. ESI-MS (pos): m/z 436.2 $[M+H]^+$

**Synthesis of methyl 4-(octyloxy)benzoate.**

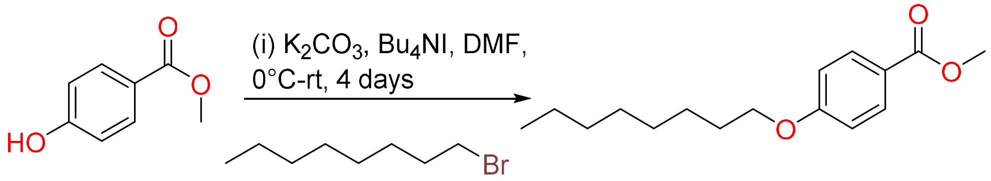


To a stirred solution of 4-hydroxy methyl benzoate (2.0 g, 13.1 mmol) was dissolved in dry DMF (30 mL) was added $K_2CO_3$ (5.43 g, 39.3 mmol), $Bu_4NI$ (4.83 g, 13.1 mmol), and octyl bromide at ice cold water bath. The reaction was allowed to room temperature. After the reaction mixture stirred at rt for 96 h, the reaction mixture was quenched with water (50 mL) and extracted into ethyl acetate (40 mL x 3). The combined organic layers were washed with brine and dried over anhydrous $Na_2SO_4$. The crude mixture was purified by silica gel column chromatography to obtain the pure product 3 (0.7 g, yield 40%, based on recovered starting material, BRSM). $^{1}$H-NMR (400 MHz, $CDCl_3$) δ 7.90 (d, *J* = 8.9 Hz, 2H), 6.83 (d, *J* = 8.9 Hz, 2H), 3.93 (t, *J* = 6.6 Hz, 2H), 3.81 (s, 3H), 1.72 (dt, *J* = 14.7, 6.7 Hz, 2H), 1.37 (q, *J* = 7.5 Hz, 2H), 1.33-1.15 (m, 8H), 0.89-0.74 (m, 3H). $^{13}$C-NMR (101 MHz, $CDCl_3$) δ 166.9, 163.0, 131.6, 122.3, 114.1, 68.2, 51.8, 31.8, 29.3, 29.2, 29.1, 26.0, 22.7, 14.1.

**Synthesis of 4-(octyloxy)benzoic acid.**

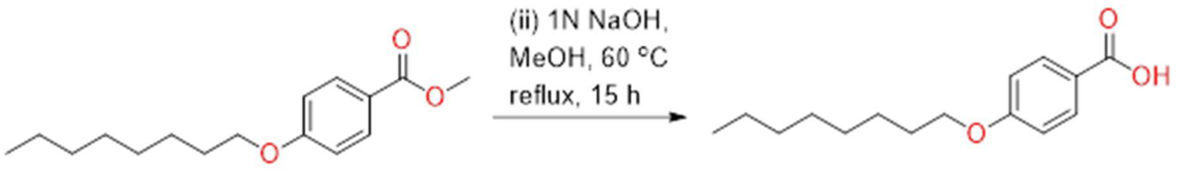


To a solution of methyl ester (0.7 g, 2.64 mmol) in MeOH (12 mL), was slowly added a aqueous solution of 1N NaOH (3 mL) and the reaction was heated to 60 °C for 15 h. Upon completion of the reaction by TLC, solvents were removed under reduced pressure. The residue was redissolved in ethyl acetate (60 mL), and the pH was adjusted to 3-4 using 3N aqueous HCl. The aqueous layer was extracted with EtOAc (10 mL×2) and the combined EtOAc layers were washed with water (1o mL) and dried over anhydrous $Na_2SO_4$. EtOAc was removed under reduced pressure, and the crude was purified by $SiO_2$ flash chromatography using (0-70% EtOAc-hexanes) to afford the corresponding acid as a white solid 4 (0.65 g, yield 98%). $^{1}$H-NMR (400 MHz, $CDCl_3$) δ 7.98 (d, *J* = 8.9 Hz, 2H), 6.86 (d, *J* = 8.9 Hz, 2H), 3.95 (t, *J* = 6.6 Hz, 2H), 1.81-1.64 (m, 2H), 1.39 (td, *J* = 8.5, 4.5 Hz, 2H), 1.33-1.14 (m, 8H), 0.87-0.74 (m, 3H). $^{13}$C-NMR (101 MHz, $CDCl_3$) δ 171.7, 163.7, 132.4, 121.3, 114.2, 68.3, 31.8, 29.3, 29.2, 29.1, 26.0, 22.7, 14.1.

**2-fluoro-6-((4-(heptyloxy)phenyl)sulfonamido)-3-methylbenzoic acid, 7.**

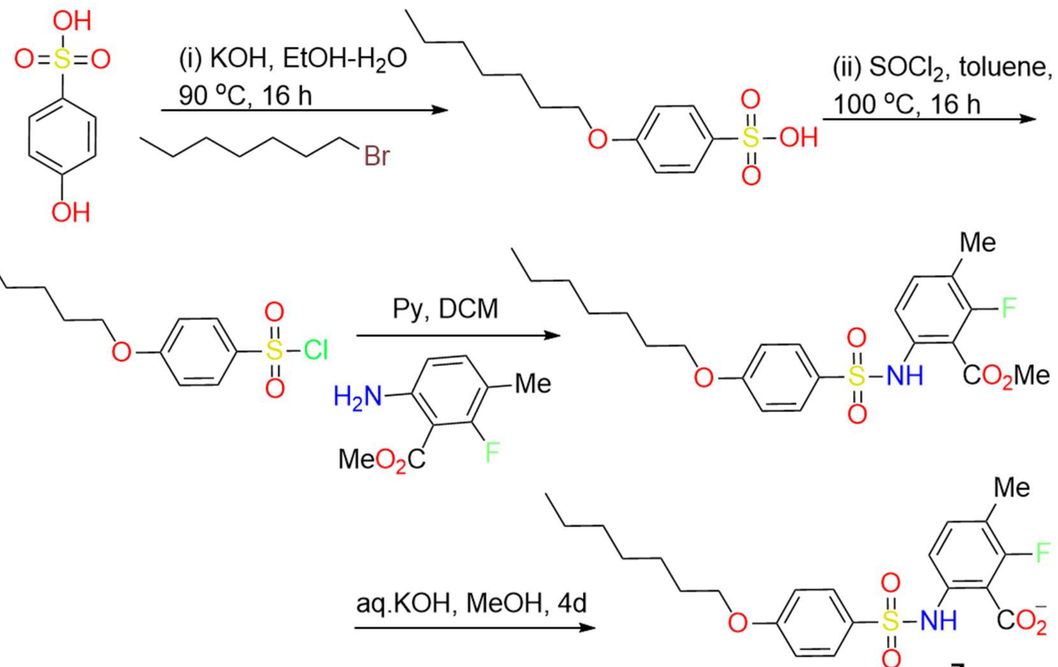


**Synthesis of 2-fluoro-6-[(4-heptoxyphenyl)sulfonylamino]-3-methyl-benzoic acid, 7.**

To a stirred solution of methyl 2-fluoro-6-[4-heptoxyphenyl)sulfonylamino]-3-methyl-benzoate (198 mg, 0.453 mmol) and KOH (50.8 mg, 0.905 mmol) in THF, Water (1:1) and stirred at 60 °C. After 4 days, the solvent was removed under reduced pressure. Reaction mixture was redissolved in minimum amount of water, acidified with 1M HCl. The precipitated product was filtered. Filtrate was washed with ethyl acetate (20 mL x 3), combined organic layers were washed with saturated brine and dried over anhydrous $Na_2SO_4$. The solvent was removed under reduced pressure. The crude product was purified by silica gel column chromatography using hexane-ethyl acetate solvent as a mobile phase to provide the desired product **7**. $^1$H-NMR (400 MHz, MeOH-d4) δ 7.60-7.63 (m, 2H), 7.30 (t, *J* = 4.0 Hz, 2H), 6.92-6-94 (m, 2H), 3.98 (t, *J* = 12.8 Hz, 2H), 2.18 (s, 3H), 1.73-1.76 (m, 2H), 1.29-1.44 (m, 8H), 0.88-0.91 (m, 3H). $^{19}$F-NMR (376.3 MHz, DMSO) δ -5.95. ESI-MS (pos): m/z 446.1 $[M+Na]^+$

**Synthesis of methyl 6-amino-2-fluoro-3-methyl-benzoate.**

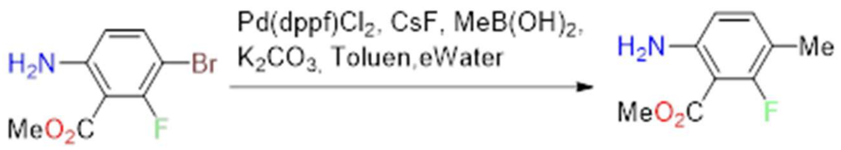


Methyl 6-amino-3-bromo-2-fluoro-benzoate (1.00 g, 4.03 mmol, combi-blocks, Cat. no.: HD-1855; CAS no. 1637774-20-9/1g/$370), Pd(dppf)$Cl_2$ (0.295 g, 0.403 mmol), CsF (0.735 g, 4.84 mmol), $MeB(OH)_2$ (0.483 g, 8.06 mmol), $K_2CO_3$ (1.67 g, 12.1 mmol) were combined in a MW vessel and were dissolved in 5:1 Toluene/Water. The reaction was then allowed to heat in the microwave reactor for 15 hours at 110 °C. LCMS analysis showed that the product was present and correlated to the main UV peak. After removal of solvent under reduced pressure, the sample was purified using a 10g NP-Flash column using hexane and ethyl acetate as the mobile phase to furnish the desired product, which can be used directly for the next step.

**Synthesis of 4-heptoxybenzenesulfonate.**

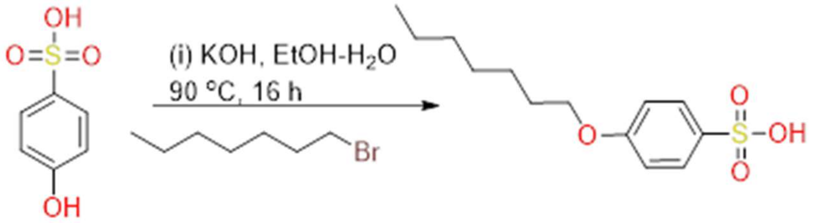


To a stirred solution of 1-bromoheptane (9089 μL, 57.8 mmol), 4-hydroxybenzenesulfonate (9100 mg, 52.6 mmol), KOH (3243 mg, 57.8 mmol) in Ethanol and water, was refluxed at 90 °C overnight. After 16 hours, another 1700 mg of KOH (1.05 eq) was dissolved in 2.5 mL of water and was added to the reaction, the solution was then allowed to reflux for another 2 hours. The reaction was removed from heat, rotavapor and the precipitated solids were filtered. The solids were dried under vacuum, subsequently used for the next reaction without further purification.

**Methyl 2-fluoro-6-[(4-heptoxyphenyl)sulfonylamino]-3-methyl-benzoate.**

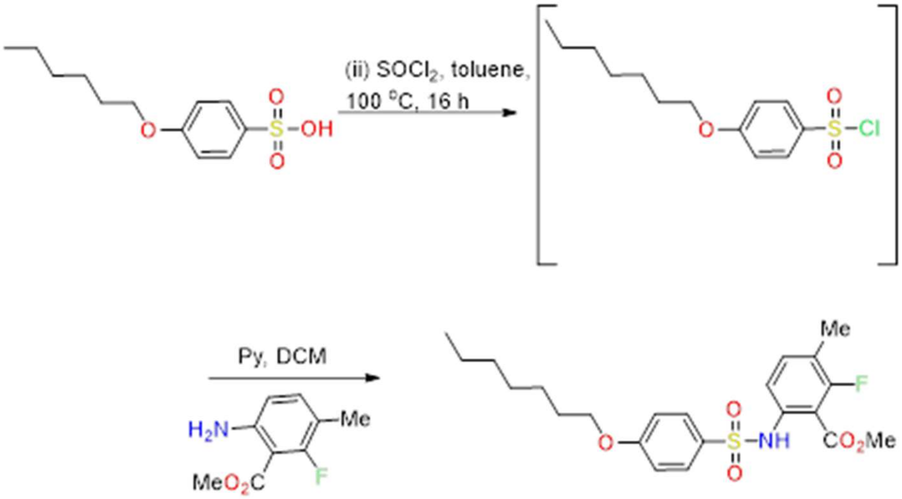


4-heptoxybenzenesulfonate (6500 mg, 24.0 mmol) was dissolved in anhydrous toluene in an RB and then the $SOCl_2$ (8736 μL, 120 mmol) was added dropwise. The reaction was allowed to reflux at 100 °C overnight. The sample was rotavapor to dryness and redissolved in dry DCM, subsequently added to a solution containing methyl 6-amino-2-fluoro-3-methyl-benzoate (742 mg, 4.05 mmol) in anhydrous DCM and pyridine (894 mg, 11.3 mmol) dropwise at RT. After 2 hours, the reaction mixture was concentrated to remove the solvent, the crude mixture was purified by silica gel column purification using hexane and ethyl acetate as a mobile phase to obtain the desired methyl ester product which was directly used for the next step.

MC1

### Abbapolins Suppress 22Rv1 Tumor Growth in Mice

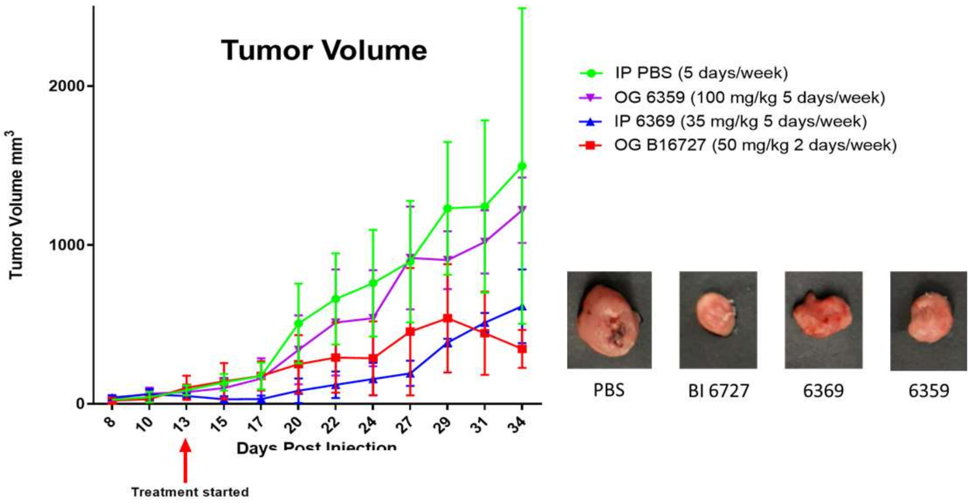


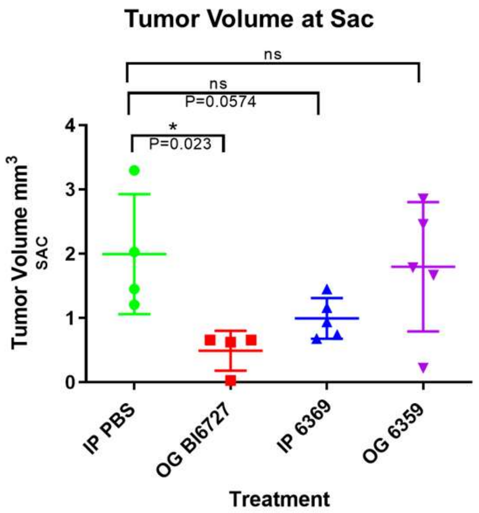


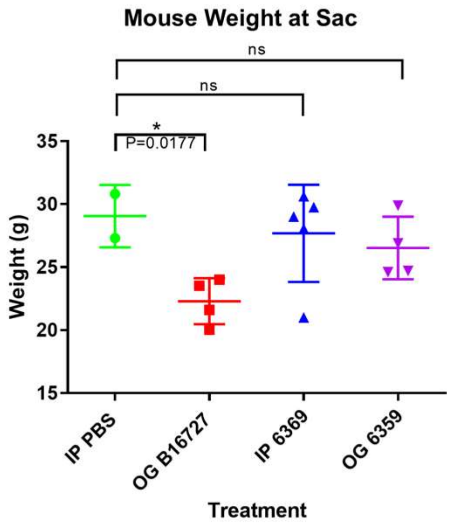


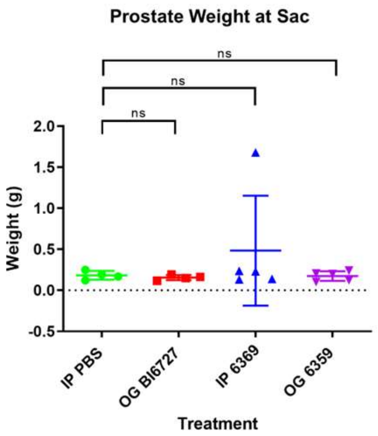


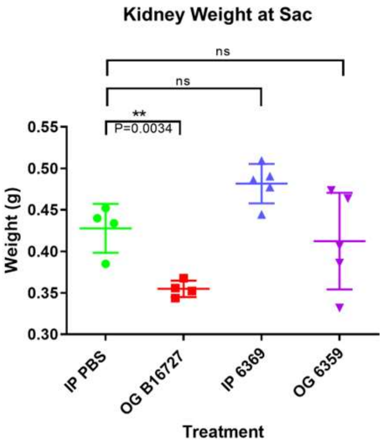


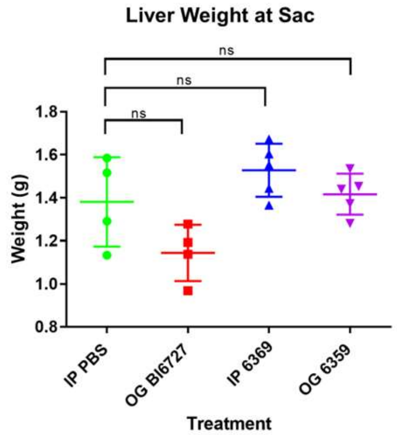


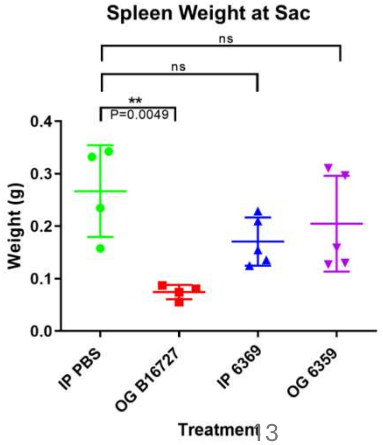


Supplementary Figure 1



In Collaboration with the Pena lab, Dep. Of Biological Sciences, UofSC

#### Slide 13

**MC1** Abbapolins suppress 22RV1.....

Mcinnes, Campbell, 2022-10-24T18:38:19.358

### Abbapolins Suppress PC3 Tumor Growth in Mice

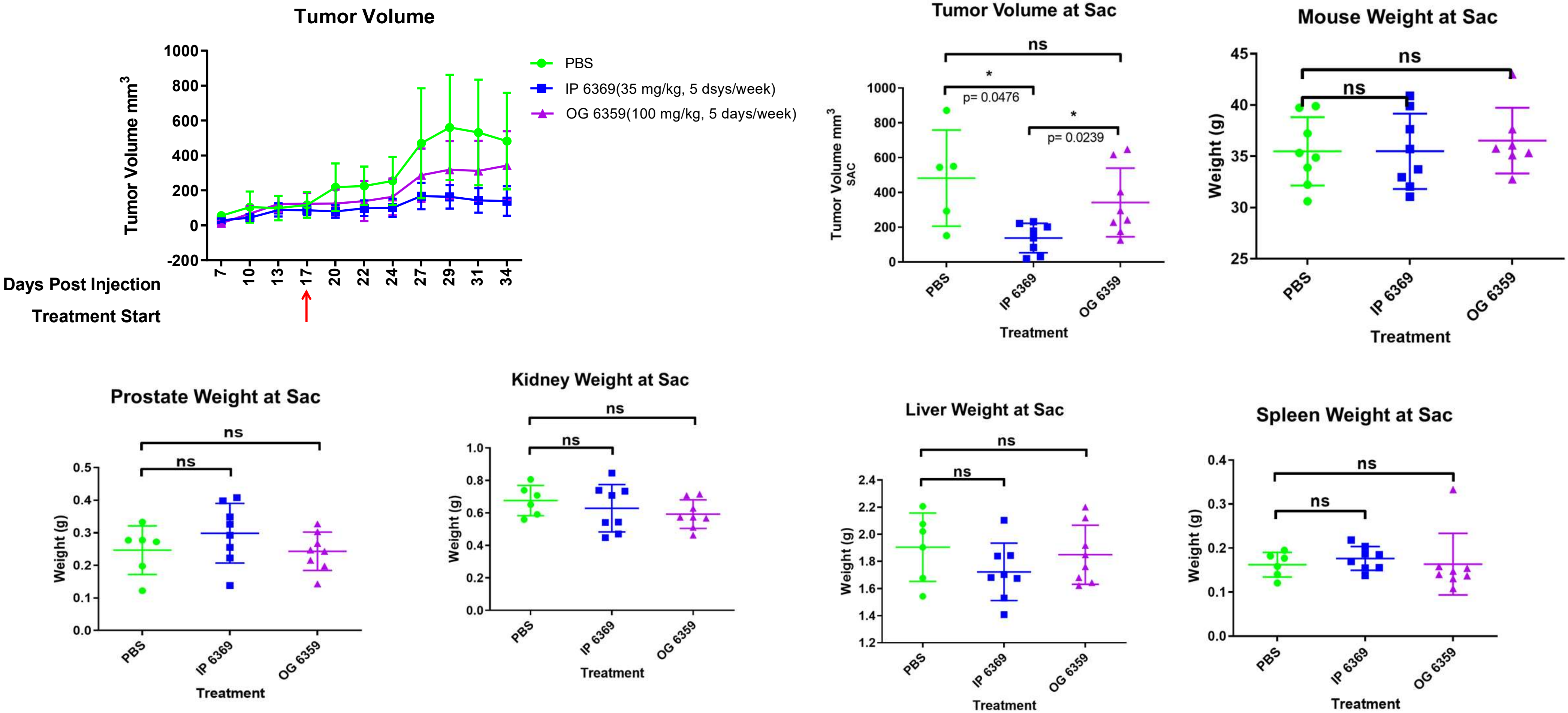


Supplementary Figure 2



In Collaboration with the Pena lab, Dep. Of Biological Sciences, UofSC
